# Human HLF^neg^ placental erythro-myeloid progenitors give rise to HLA Class II^neg^ Hofbauer cells

**DOI:** 10.1101/2022.02.26.482080

**Authors:** Jake R. Thomas, Anna Appios, Emily F. Calderbank, Xiaohui Zhao, Russell S. Hamilton, Ashley Moffett, Andrew Sharkey, Elisa Laurenti, Naomi McGovern

## Abstract

The earliest macrophages are generated during embryonic development from erythro-myeloid progenitors (EMPs) via primitive haematopoiesis. This process is still poorly understood in humans but is generally thought to be spatially restricted to the yolk sac. Human fetal placental macrophages, Hofbauer cells (HBC), arise during the primitive haematopoietic wave, yet are unlikely to be yolk sac derived as they appear prior to placental vascularisation. Here we identify a population of placental erythro-myeloid progenitors (PEMPs) in the early human placenta that give rise to HBC. PEMP are fetal CD34^+^CD43^+^ progenitors found exclusively at early gestational timepoints. Transcriptomic analyses reveal that PEMP have a unique transcriptome with some conserved features of primitive yolk sac EMPs, including the lack of *HLF* expression. Using *in vitro* single-cell culture experiments we show that PEMP generate HBC-like cells which lack HLA-DR expression, a conserved feature of all fetal primitive macrophages in humans. These findings indicate that HBC are derived locally from PEMP and demonstrate that human primitive haematopoiesis is not restricted to the yolk sac, occurring also in the placenta.

## Introduction

Tissue-resident macrophages (TRMs) appear early in gestation and are likely to have critical roles in organogenesis and fetal development^1, 2^. The first TRMs are generated by primitive haematopoiesis from a population of ‘early’ erythro-myeloid progenitors (EMPs) in the yolk sac^3–5^. Later in gestation transient definitive “late” EMPs and definitive haematopoietic stem cells (HSCs) give rise to TRMs via a monocyte stage in many tissues^4, 6–10^. Primitive haematopoiesis has been studied in mice^4, 11, 12^, but our understanding remains limited in humans^13^, because it is currently impossible to distinguish between primitive and definitive haematopoietic progeny.

Human placental TRMs, Hofbauer cells (HBC), are likely to be primitive TRMs as they are present from day 18 of gestation^14^ (prior to the vascularisation of the placental mesenchyme^15^), and are transcriptionally similar to yolk sac macrophages^16^. Since fetal blood flow to the placenta is not established until the ∼10^th^ week of gestation^17, 18^, it is likely that primitive HBC are derived from local progenitors.

In this study we show that human primitive macrophages can be distinguished from definitive macrophages by their expression of HLA Class II molecules. This finding is consistent across a range of human fetal macrophages, including HBC. We subsequently identify a population of placental erythro-myeloid progenitors (PEMP) with features of primitive haematopoietic progenitors. We go on to show that PEMP can differentiate into primitive HLA-DR^neg^ HBC-like macrophages *in vitro*. These findings provide significant additional insight into the nature of human primitive macrophages and confirm that the placenta is a site of primitive haematopoiesis.

## Results

### HLA Class II expression delineates primitive and definitive human macrophages across tissues

We have previously shown that a shared feature of first trimester HBC and the earliest yolk sac macrophages is a complete lack of expression of HLA class II molecules (**Fig.S1A**)^16^. To see if this is conserved across all primitive macrophages we analysed publicly available scRNAseq gene expression data from a range of early human embryonic samples - Carnegie stage 7 to 11 (CS7 to CS11)^19–21^. Combined analysis reveals a population of macrophages conserved across all datasets (**Fig.1A,B, Fig.S1B,C**) that express prototypical macrophage genes, including *PTPRC*, *CD68*, *CD14*, *AGR2*, *CD36* and *AIF1* (**Fig.1C**), and that uniformly lack the expression of HLA class II genes (*HLA-DRA, HLA-DRB1, HLA-DRB5, HLA-DQA1, HLA-DQB1*) (**Fig.1C**).

**Fig. 1.**
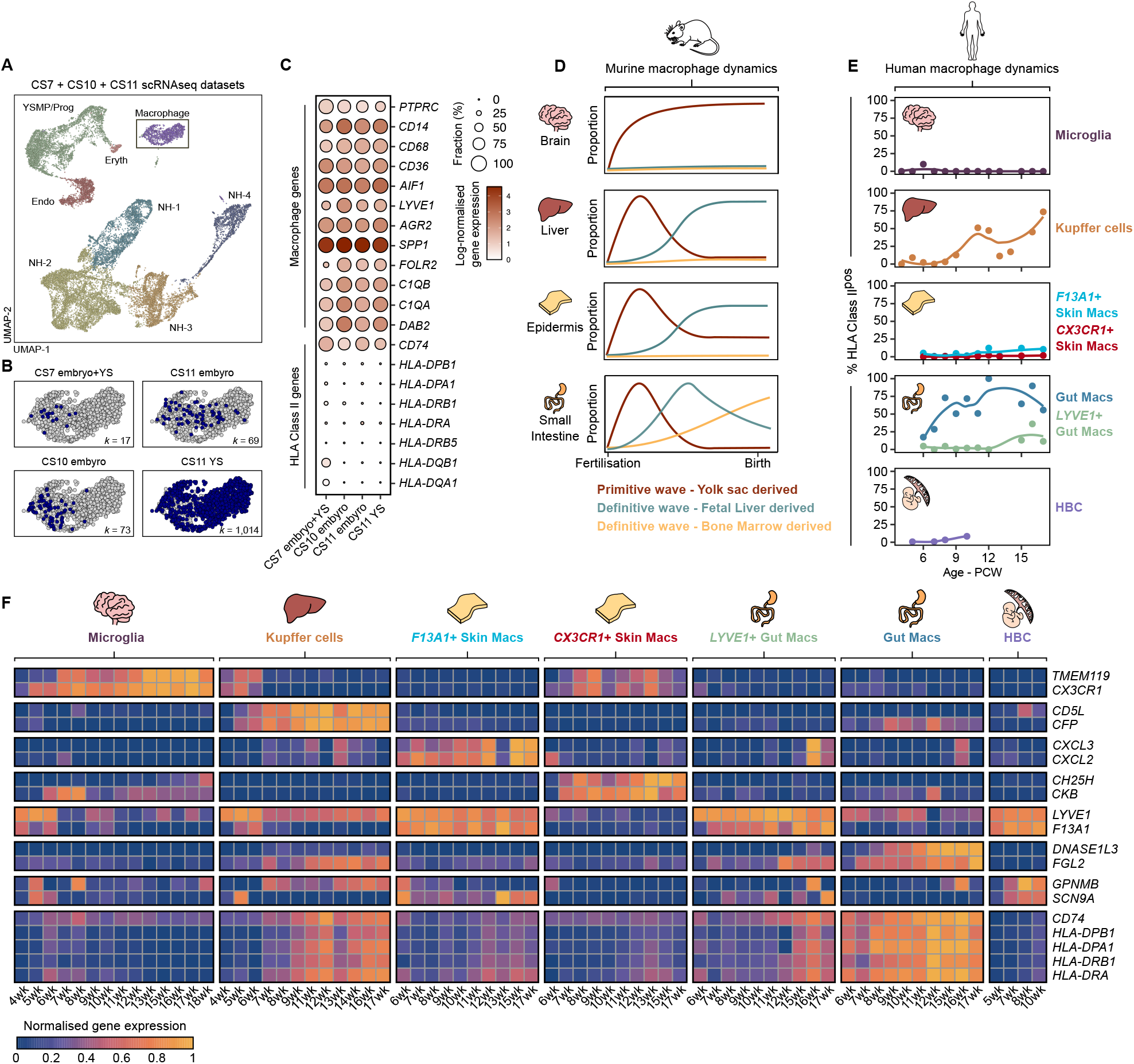
Human primitive macrophages are HLA-DR^neg^. **A**) UMAP visualisation of 18,655 human early embryo single-cell transcriptomes from^19–21^, NH– Non-haematopoietic, Endo – Endothelial, YSMP – Yolk sac myeloid-biased progenitors, Prog – Haematopoietic progenitors, Eryth – Erythrocytes. **B**) UMAP visualisation of early embryonic macrophage single-cell transcriptomes, cells from each study are highlighted in blue within each panel. Number of cells from each dataset (*k*) are annotated on each panel. CS – Carnegie Stage, YS – Yolk sac. **C**) Dotplot heatmap displaying log-normalised gene expression of genes associated with macrophage identity and HLA Class II expression, within early human macrophages. Dot size represents fraction of cells with nonzero expression. **D**) Diagram depicting the previously established contributions of primitive and definitive haematopoiesis to tissue-resident macrophage populations across murine organs. Redrawn and modified from^22^. **E**) Line graphs showing the temporal dynamics of HLA Class II expression in tissue-resident macrophages (TRMs) from different human fetal tissues^20, 23–27^. Cells are considered HLA Class II^pos^ if they display non-zero expression of *HLA-DRA*, *HLA-DRB1*, *HLA-DPA1*, *HLA-DPB1*, *HLA-DQA1* and *HLA-DQB1*. HBC – Hofbauer cells, PCW – Post-conception weeks. **F**) Normalised gene expression heatmap of marker genes and HLA Class II genes in human fetal TRMs across developmental time.

Fate-mapping models by others have established macrophage turnover in murine tissues during fetal development (**Fig.1D**)^5–8, 22^. We propose that the dynamics of gene expression in fetal TRMs, in particular HLA class II expression, can be used to visualise macrophage turnover in humans in lieu of fate-mapping tools (**Fig.1E,F Fig.S2A-D**)^20, 23–27^. We find macrophages adopt tissue-specific gene signatures in a time dependent fashion, for example *TMEM119* and *CX3CR1* in microglia and *CD5L* and *CFP* in Kupffer cells (**Fig.1F**). The temporal expression dynamics of HLA class II genes in human fetal TRMs map closely to murine macrophage ontogenies established by fate-mapping models(**Fig.1D-F**)^22^. For example, microglia, which have minimal contributions from definitive haematopoiesis^5, 6^, remained negative for HLA class II up to 18 Post-Conception weeks (PCW) (**Fig.1E,F, Fig.S2D**). In contrast, Kupffer cells, where primitive cells are replaced by fetal liver-derived definitive cells in mice^6^, are HLA class II-negative at early timepoints (4-7 PCW) but positive later (11wk-17wk PCW) (**Fig.1E,F, Fig.S2D**).

Therefore, a conserved feature of human primitive macrophages in embryonic tissues is the lack of expression of HLA class II genes, with the expression of these genes providing insight into macrophage replacement dynamics across fetal tissues.

### Placental erythro-myeloid progenitors (PEMP) are only present in early gestation

The observation that HBC are present 18 days post fertilisation (dpf)^14^, before fetal blood flow to the placenta begins suggests that they arise from local progenitors. Indeed, CD43^+^CD34^+^ progenitors have been described in the first trimester placenta but are poorly defined^15, 28, 29^. Due to the early appearance of HBC and the lack of fetal monocytes in the first trimester placenta^16^, we propose that these CD43^+^CD34^+^ progenitors are placental EMPs (PEMP). To develop our understanding of CD43^+^CD34^+^ placental PEMP, we designed a flow cytometry panel to isolate these cells, excluding any lymphocyte, erythrocyte, monocyte/macrophage and granulocyte contamination (**Fig.2A**). HLA-DR^-^FOLR2^-^CD41^-^Lin^-^CD66b^-^CD14^-^CD235a^-^ CD34^+^CD43^+^ PEMP express haematopoietic makers CD45 and c-KIT (CD117), but lack expression of CD90 (expressed by definitive HSCs in the bone marrow^30^ and placental fibroblasts) (**Fig.2B**), EGFR and HLA-G (trophoblast markers)^31^ (**Fig.S3A**). Their fetal origin is confirmed by staining with HLA-allotype antibodies (**Fig.2C**) and typical haematopoietic progenitor morphology is seen in Geimsa-stained cytospins of FACS-sorted PEMP (**Fig.2D, Fig.S3B**). Although rare, CD43^+^ CD34^+^ CD45^+^ PEMP are reliably observed within the stroma of first trimester placental villi by immunofluorescence. We find PEMP in close association with CD34^+^ placental endothelium(**Fig.2E, Fig.S3C**), in line with previous reports of EMPs arising from endothelial cells^32^. Applying our flow cytometry gating strategy to placental samples of different gestational ages reveals that PEMP decrease significantly after the first weeks of pregnancy, and they are virtually absent from 9 weeks estimated gestational age (EGA) (**Fig.2F-H**). The ratio of HBC:PEMP increases over gestation, consistent with PEMP initially seeding a minor population of macrophages which then proliferate and expand *in situ* (**Fig.S3D-E**).

**Fig. 2.**
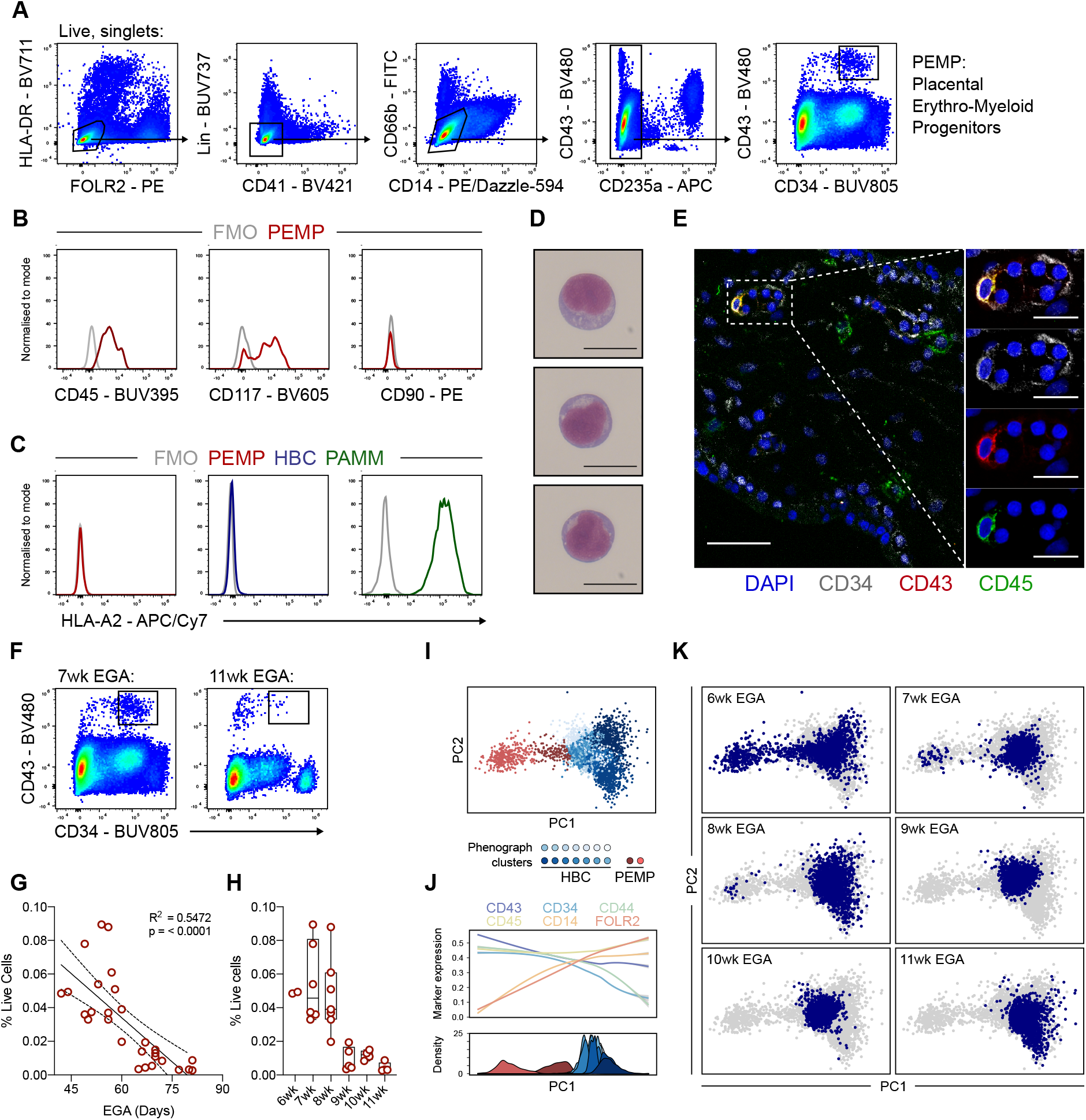
Placental erythro-myeloid progenitors (PEMP) are exclusively found early in gestation. **A**) Representative flow cytometry gating strategy identifying CD43^+^ CD34^+^ placental erythro-myeloid progenitors (PEMP) in a 7wk EGA (estimated gestational age) sample. **B**) Flow cytometric expression of CD45, CD117 (c-Kit) and CD90 in PEMP, data shown are representative of *n* = ≥2. FMO – fluorescence minus one. **C**) Relative flow cytometry expression of HLA-A2 in PEMP, HBC (Hofbauer cells) and PAMM (Placenta-associated maternal macrophages) from a sample with an HLA-A2 allotype mismatch between mother and fetus. **D**) Representative Giemsa-stained cytospins of PEMP isolated by FACS. Scale bars, 20μm. Representative images from *n* = 3 donors. **E**) Identification of CD34^+^ (grey) CD43^+^ (red) CD45^+^ (green) PEMP within the villous stroma of a 7wk EGA placental sample. Scale bars, main panel 100μm, inset panels 20μm. **F**) Flow cytometric analysis of PEMP abundance in two samples of distinct EGA. **G**) Quantification of PEMP abundance via flow cytometry as a proportion of all live cells, plotted against EGA by day, or **H**) grouped by week. (*n* = 28). **I**) PCA (principle component analysis) embedding of flow cytometry data of HBC, PEMP and any phenotypic intermediates from 12 placental samples (gating strategy for analysis detailed in Fig.S3.F,G). Cells were subjected to Phenograph clustering and clusters are coloured according to their cellular identities. (*n* = 12, 2x 6wk EGA, 2x 7wk EGA, 2x 8wk EGA, 2x 9wk EGA, 2x 10wk EGA and 2x 11wk EGA samples). **J**) Mean expression of key flow cytometric markers (top panel), and distributions of phenograph clusters (bottom panel) along PC1. **K**) PCA embeddings as in (I), with cells from each timepoint highlighted in blue within each panel.

UMAP visualisation of flow cytometry data from a 6wk EGA sample reveals some PEMP-HBC intermediates might be missed by our gating strategy (**Fig.S3F)**. To further investigate this population, we used Hyperfinder to derive a gating strategy to isolate HBC, PEMP and any potential phenotypic intermediates (**Fig.S3F-I**). We applied this gating strategy to 2 donors from each EGA group, generated a principle component analysis (PCA) embedding and performed Phenograph clustering on the combined dataset (**Fig.S3J, Fig.2I**). Proposed PEMP-HBC differentiation is captured along the first principle component (PC1), and analysis of marker expression reveals one cluster of PEMP with elevated expression levels of CD14 and FOLR2 indicating cells are in transition from PEMP to HBC (**Fig.2J, Fig.S3I**). The cells are only seen at 6wk EGA whilst PEMP are found at 6-8wk EGA (**Fig.2K**), suggesting that potential PEMP-HBC differentiation only occurs at earlier timepoints.

These findings demonstrate that the human placenta contains a transitory population of fetal haematopoietic progenitors, PEMP, which are found only at early timepoints in gestation.

### Transcriptomic and phenotypic heterogeneity of PEMP

To confirm that PEMP are true haematopoietic progenitors we first looked for haematopoietic progenitors in a scRNAseq atlas of the first trimester placenta^26^. Integration of this dataset with other fetal scRNAseq datasets (CS7 embryo^19^, CS10 embryo^21^, CS11 YS^20^, CS15 aorta-gonad mesephros (AGM)^21^, fetal liver^23^ and fetal bone marrow (BM)^33^) reveals very few placental cells in the combined haematopoietic progenitor cluster (**Fig.S4A**). This is likely due to the scarcity of PEMP and their presence only early in gestation. Therefore, we performed SmartSeq2 scRNAseq on PEMP, HBC, an undefined population of CD43^+^CD34^-^ cells, and endothelial cells from a 6wk and a 7wk EGA sample (**Fig.S4B**). After pre-processing, and the removal of 35 endothelial cells (*CDH5*+, *KDR*+) and 10 erythrocytes (*HBZ*+, *HBE1*+) from the dataset, 210 cells were annotated into 6 clusters based on their gene expression (**Fig.3A**). All cells profiled are fetal in origin, and each cluster contains cells derived from each donor (**Fig.S4C,D**). The populations identified are: 3 PEMP (PEMP-1,2,3), HBC, megakaryocyte progenitors (Mega), and a population composed of both PEMP and HBC (PEMP-HBC). Cells within the PEMP clusters are primarily derived from the PEMP gate during FACS isolation (**Fig.S4E**). Analysis of differentially expressed genes (DEGs) between the clusters reveals heterogeneity within the PEMP compartment, including cell surface markers and transcription factors (**Fig.S5A-D**). PEMP-1 express typical haematopoietic progenitor genes including *HOXA9*, *CD34*, *SPINK2*, *HOPX* (**Fig.3B**). PEMP-2 and PEMP-3 are myeloid and Mega/Eryth/Mast (MEM) biased progenitors respectively, based on their expression of known lineage-specific genes *IRF8*, *MPO*, *CLEC11A CSF3R*, *GATA1*, *GATA2* and *KIT* (**Fig.3B**). HBC and Mega clusters express known macrophage (*C1QA*, *CD14*, *CD163*) and megakaryocyte (*THBS1*, *PF4*, *ITGA2B*) genes (**Fig.3B**).

**Fig. 3.**
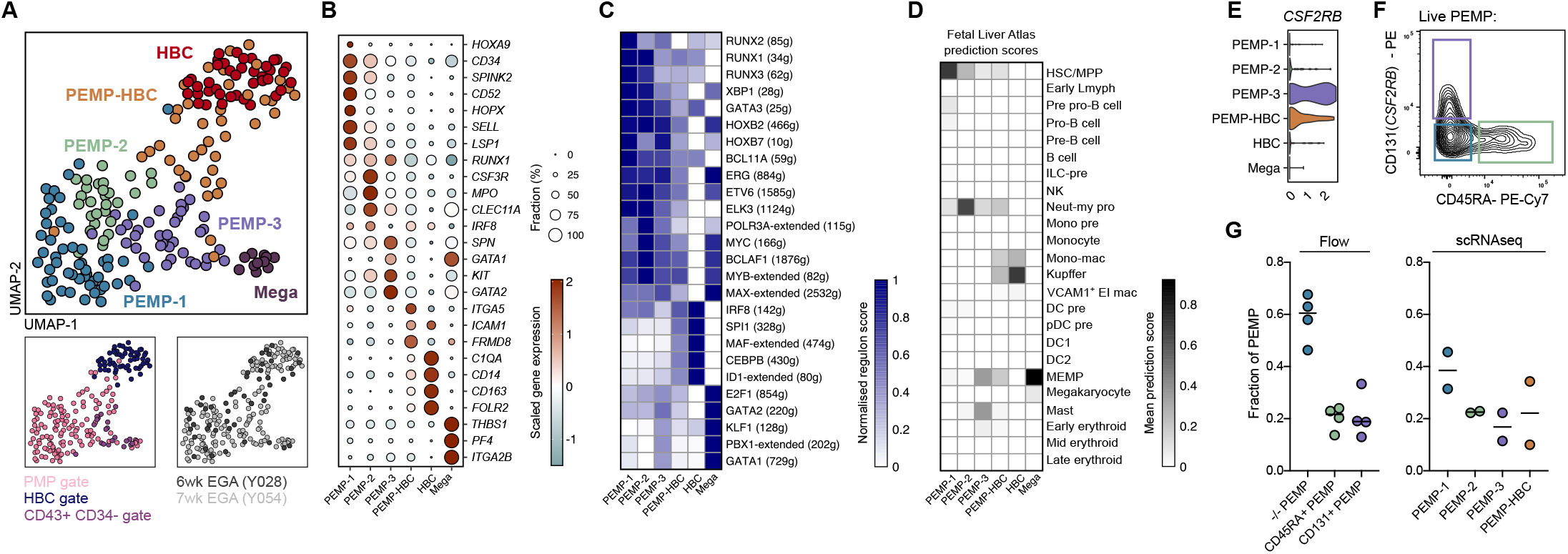
PEMP display transcriptomic and phenotypic diversity. **A**) UMAP visualisation of 210 single-cell transcriptomes from 2 early EGA placental samples. Cells are coloured by cluster identity. Bottom panels, cells are coloured by gate used for FACS isolation (left) and by donor (right). Mega – Megakaryocyte progenitors. **B**) Dotplot heatmap displaying scaled log-normalised gene expression of key marker genes across the observed clusters. Dot size represents fraction of cells with nonzero expression. **C**) Heatmap of selected inferred transcription factor activities (regulon scores) across the observed clusters, calculated via SCENIC analysis^34^. The number of genes associated with each regulon is listed in parentheses. **D**) Heatmap showing mean prediction scores (transcriptomic similarity) of single cells within observed placental clusters using a human fetal liver atlas dataset as a reference^23^. Scores were calculated using the FindTransferAnchors function in Seurat (see Methods). **E**) Violin plot of *CSF2RB* log-normalised gene expression across observed clusters. **F**) Analysis of CD131 and CD45RA expression within PEMP via flow cytometry revealing three populations: ^-/-^ PEMP (blue), CD45RA^+^ PEMP (green) and CD131^+^ PEMP (purple). **G**) Quantification of PEMP populations via flow cytometry (*n*=4) (left panel), and scRNAseq clusters within the PEMP gate (right panel).

To further probe PEMP identity we leveraged SCENIC^34^ to infer transcription factor (regulon) activities within each cluster (**Fig.S5E, F)**. PEMP are under the control of transcription factors important in haematopoietic progenitor identity and fate, including RUNX1, RUNX3, GATA3 and ERG (**Fig.3C**). In line with our gene expression data, PEMP-2 and PEMP-3 express transcription factors typical of myeloid lineages (MYC, MYB, IRF8) and MEM lineages (GATA1, GATA2, KLF1, PBX1) respectively (**Fig.3C**).

Using the *FindTransferAnchors* function in Seurat we analysed the transcriptomic similarities between our clusters and annotated clusters from a recently published dataset of human fetal liver haematopoiesis^23^. Each PEMP cluster displays similarity to distinct fetal liver progenitors; PEMP-1 to HSC/MPP, PEMP-2 to Neut-my Pro (granulo-myeloid progenitors - GMPs) and PEMP-3 to MEMP (**Fig.3D**). A minor population of cells within the PEMP-3 cluster display high transcriptomic similarity to mast cells (**Fig.3D, S6A**). These cells express canonical mast cell genes (*HDC*, *MITF*, *CPA3*, *KIT*), but lack the expression of *FCER1A* mRNA (**Fig.S6B**). Flow cytometry analysis confirms the presence of fetal CD200R^+^CD117^+^FCεR1^-^ mast cells in early placental samples (**Fig. S6B**). Placental mast cell FCεR1 expression is consistent with previous reports describing fetal mast cells^35, 36^ but further investigation is beyond the scope of this study.

These analyses suggest that PEMP-1 can differentiate along two distinct pathways governed by unique transcriptional programmes: either towards the myeloid lineage (PEMP-2) or towards the MEMP lineage (PEMP-3). CD131 (*CSF2RB*) is expressed specifically in PEMP-3 (**Fig.3E, S5B**), and CD45RA is known to be expressed by definitive GMPs within the adult bone marrow^37^. We can identify three PEMP populations (CD45RA^+^ PEMP, CD131^+^ PEMP and ^-/-^PEMP) via flow cytometry (**Fig.3F, Fig.S6D, E**) and find they are similarly abundant to the clusters in our scRNAseq data (**Fig.3G**), validating our observed transcriptional heterogeneity.

In summary, our findings confirm that PEMP are true haematopoietic progenitors, and suggest that PEMP seed diverse haematopoietic populations within the placenta.

### PEMP display characteristics of primitive erythro-myeloid progenitors and do not express *HLF*

We predict that PEMP are primitive progenitors because of their temporal distributions throughout gestation, and the expression status of HLA Class II in HBC, their proposed progeny. We therefore compared their transcriptomes with other populations of human fetal haematopoietic progenitors (**Fig.4A, S7A,B**)^19–21, 33^. Hierarchical clustering of the integrated dataset reveals a broad division between the progenitor populations, with PEMP-1 clustering most closely to primitive CS7 EMPs, away from yolk sac myeloid-biased progenitors (YSMPs), aorta gonad mesonephros (AGM), fetal liver (FL) and bone marrow (BM) HSCs (**Fig.4B**). The clustering of YSMP with HSCs suggests that these cells are definitive progenitors. To determine conserved features of primitive EMP we calculated differentially expressed genes between both CS7 EMPs and PEMP-1 and definitive progenitors (**Fig.4C**). *HLF* is not expressed in either CS7 EMPs and PEMP, but is present in all other lineages, including YSMP (**Fig.4D**). *HLF* is known to be constitutively expressed in murine definitive HSC, but not EMPs^38^.

**Fig. 4.**
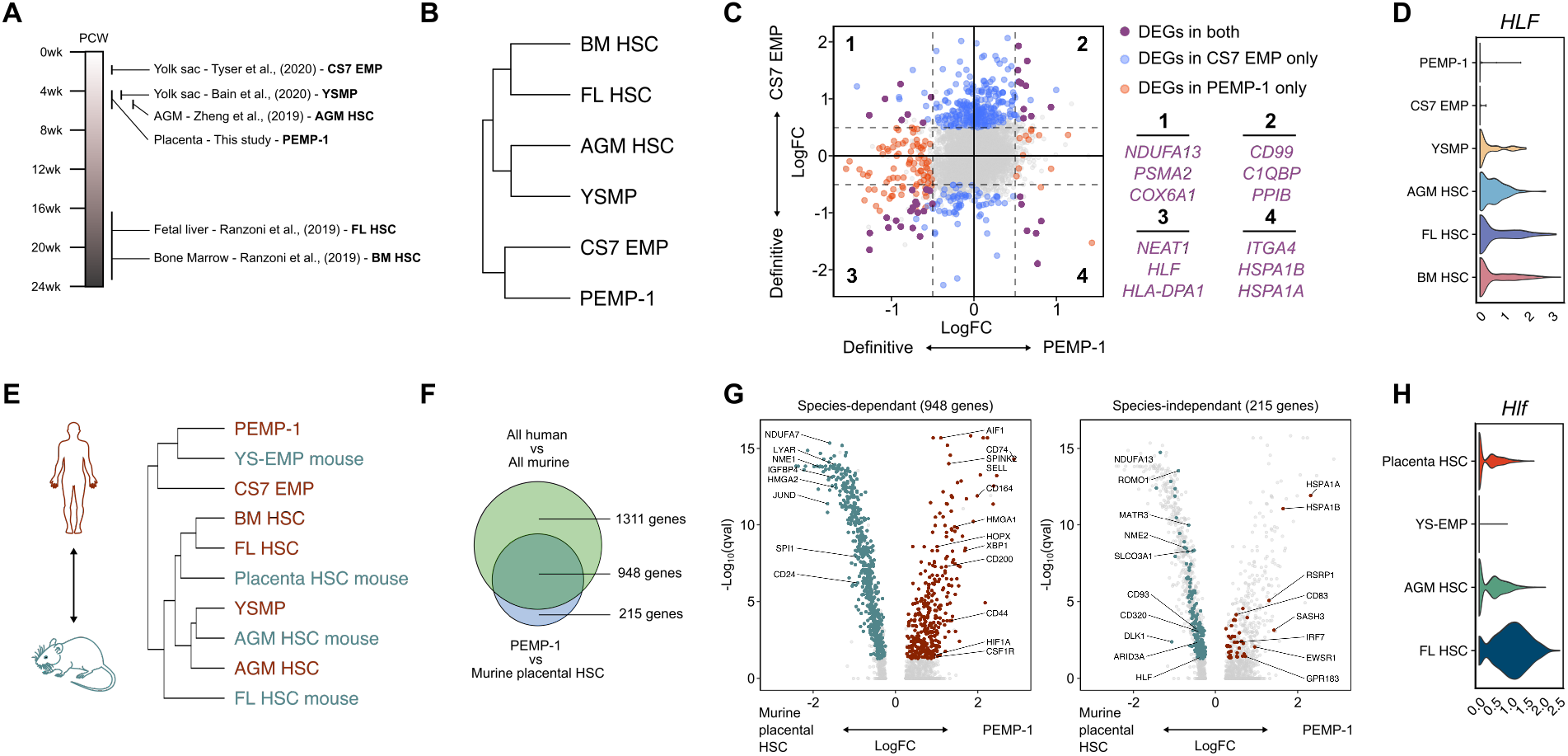
PEMP display characteristics of primitive erythro-myeloid progenitors. **A**) Schematic representation of human haematopoietic progenitor populations used for comparative analysis^19–21, 33^. CS – Carnegie Stage, EMP – Erythro-myeloid progenitors, YSMP – Yolk sac myeloid-biased progenitors, AGM – Aorta-gonad mesonephros, FL – Fetal liver, BM – Bone marrow, HSC – Haematopoietic stem cell. **B**) Hierarchical clustering dendrogram depicting transcriptomic similarity between human fetal progenitor populations, calculated using gene expression distance matrix from the integrated dataset (Fig.S7A). **C**) Scatter plot showing differentially expressed genes (DEGs) between PEMP-1 and definitive progenitors (YSMP, AGM HSC, FL HSC and BM HSC) on the x-axis and CS7 EMP and definitive progenitors on the y-axis. Genes significantly differentially expressed (adjusted p value < 0.05) in both comparisons, PEMP-1 alone and CS7 EMP alone are shown in purple, orange and blue respectively. Selected genes which were differentially expressed in both CS7 EMP and PEMP-1 are highlighted. **D**) Violin plot of *HLF* log-normalised gene expression across human fetal haematopoietic progenitors. **E**) Hierarchical clustering dendrogram depicting transcriptomic similarity between human and murine^39, 40^ fetal progenitor populations, calculated using gene expression distance matrix from the integrated dataset (Fig.S7C). YS – Yolk sac. **F**) Venn diagram depicting the overlap of DEGs calculated between all human and all murine fetal haematopoietic progenitors, and DEGs calculated between PEMP-1 and murine placental HSC alone. **G**) Volcano plots showing DEGs between PEMP-1 and murine placental HSC. Left panel: genes which overlap with DEGs calculated between all human and all murine fetal haematopoietic progenitors. Right panel: genes which do not overlap. **H**) Violin plot of *Hlf* log-normalised gene expression across murine fetal haematopoietic progenitors.

Recently, the haematopoietic progenitor compartment in the murine placenta was characterised in detail^39^. To compare murine placental HSC with human PEMP we performed an inter-species transcriptional analysis (**Fig.S7C,D**). Hierarchical clustering of the integrated inter-species dataset reveals that murine placental HSC do not cluster with human primitive CS7 EMP and PEMP-1, but are transcriptionally similar to AGM, FL and BM HSCs^40^ (**Fig.4E, S7D**). We find 1,163 DEGs (adjusted p-value < 0.05) between PEMP-1 and murine placental HSC (**Fig.4F**). 948 (81.5%) are dependent on differences between species alone and are seen in other comparisons of human-murine progenitors. 215 DEGs (18.5%) are independent of differences between species comparisons (**Fig.4F,G**) and include *HLF.* Unlike PEMP-1, there is comparable *Hlf* expression between murine placental HSC and definitive AGM HSC (**Fig.4H**).

Together these data suggest that PEMP are primitive EMPs, and that the murine placental HSC that have been identified thus far are not analogous to PEMP, as they have transcriptional features of definitive progenitors.

### PEMP generate primitive HLA-DR^neg^ HBC-like cells *in vitro*

As proof-of-concept, we next explored whether PEMP can differentiate into HLA-DR^neg^ primitive macrophages. To mimic the placental microenvironment PEMP were cultured for 18 days as single cells on monolayers of primary first trimester placental fibroblasts, supplemented with cytokines that promote macrophage differentiation (**Fig.S8A**, **Fig.5A**). As a control for definitive myeloid cell differentiation, we performed identical experiments using GMPs isolated from adult peripheral blood (**Fig.S8B, Fig.5A**). The colony-forming efficiency of PEMP is significantly lower than definitive GMPs (**Fig.5B**). Despite this, PEMP generated larger colonies than GMP (**Fig.5C, Fig.S8J**), indicating they are more proliferative. PEMP single-cell cultures give rise to morphologically and phenotypically distinct colonies (**Fig.5D,E, Fig.8C-G**). They generate diverse haematopoietic lineages, including CD14^+^ cells, megakaryocytes, erythroid cells, and in rare cases granulocytes (**Fig.5D-G, Fig.S6G**), but display limited multipotency (**Fig.5F,G**). This is in contrast to YSMP, that are poor at generating erythrocytes^20^. Most colonies contain CD14^+^ cells (∼60%), and these were generated in all donors (**Fig.5G**). PEMP that give rise to CD14^+^ and megakaryocyte colonies in single-cell cultures display increased expression of CD45RA and CD131 respectively (**Fig.S8H, I**), suggesting that transcriptionally diverse populations of PEMP (**Fig.3**) have distinct fates.

**Fig. 5.**
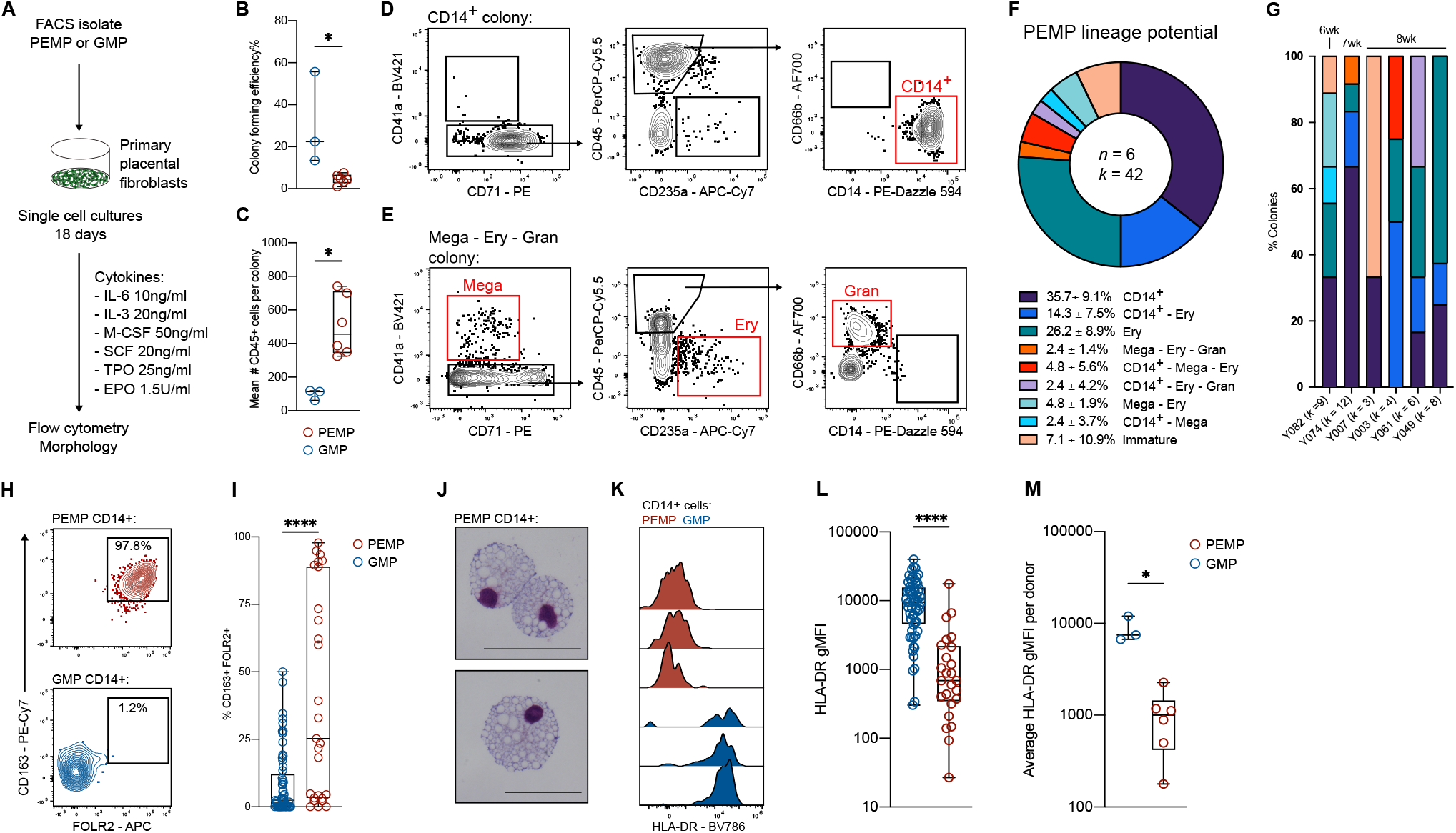
PEMP generate HLA-DR^neg^ HBC-like cells *in vitro*. **A**) Schematic depicting experimental design for PEMP single-cell culture assay. **B**) Boxplot depicting total colony forming efficiency of adult peripheral blood GMP (Granulo-myeloid progenitors) and PEMP. (GMP *n* = 3) (PEMP *n* =6). P value calculated by Mann-Whitney test. **C**) Boxplot depicting mean number of CD45^+^ cells per colony per donor. (GMP *n* = 3) (PEMP *n* =6). P value calculated by Mann-Whitney test. **D**) Representative flow cytometry analysis of a single-cell PEMP-derived colony which yielded CD14^+^ cells. **E**) Representative flow cytometry analysis of a single-cell PEMP-derived colony which yielded Megakaryocytes (Mega), erythroid cells (Ery) and granulocytes (Gran). **F**) Donut plot showing lineage potential of PEMP defined by single-cell culture and flow cytometry analysis. ‘Immature’ denotes CD45^+^ colonies which did not contain any other defined immune cell lineages. (*n* = 6 donors, *k* = 42 colonies generated). **G**) Stacked bar charts showing lineage potential of PEMP for each donor profiled. EGA of the donors are added as labels above the bar chart. **H**) Analysis of CD163 and FOLR2 expression in CD14^+^ colony derived from single PEMP and GMPs via flow cytometry, and **I**) boxplot quantification of co-expression for all CD14^+^ colonies profiled (GMP *n* = 61) (PEMP *n* = 25). P value calculated by Mann-Whitney test. **J**) Representative Giemsa-stained cytospins of cells from CD14^+^ colonies derived from a single PEMP. Scale bars, 50μm. Representative images from n=4 donors. **K**) Analysis of HLA-DR expression in CD14^+^ colony derived from single PEMPs and GMPs via flow cytometry. Each histogram represents a colony derived from an independent donor. **L**) Boxplot quantification of HLA-DR expression in CD14^+^ colony derived from single PEMPs and GMPs via flow cytometry for all colonies profiled (GMP *n* = 61) (PEMP *n* = 25). P value calculated by Mann-Whitney test. **M**) Boxplot depicting median HLA-DR expression of all CD14^+^ colonies per donor. (GMP *n* = 3) (PEMP *n* =6). P value calculated by Mann-Whitney test. *, P ≤ 0.05, **, P ≤ 0.01; ***, P ≤ 0.001, ****, P ≤ 0.0001.

To determine if PEMP-derived CD14^+^ cells are macrophages, we analysed their expression of CD163 and FOLR2, two surface receptors highly expressed by HBC^16^. PEMP seem more disposed to differentiate into CD163^+^ FOLR2^+^ macrophages (**Fig.5H**) than GMP (**Fig.5I, Fig.S8K**). PEMP-derived CD14^+^ cells are large and highly vacuolated, typical of macrophages (**Fig.5J**). Critically, PEMP-derived CD14^+^ cells have significantly less HLA-DR expression compared to GMP-derived CD14^+^ cells (**Fig.5K-M**, **Fig.S8L**). Thus, our culture system recapitulates ‘true’ primitive haematopoiesis with the generation of CD163^+^ FOLR2^+^ HLA-DR^neg^ HBC-like cells *in vitro*. In conclusion, our findings suggest PEMP found in early pregnancy produce HBC.

## Discussion

Much of our knowledge concerning embryonic macrophage generation is based on studies in mice^3–6, 12, 41^. Our work provides insight into human primitive haematopoiesis and will help future studies to shed light on human primitive macrophages. These macrophages are crucial for embryonic organ development; they aid in the remodelling of tissues via scavenger activities^16^, the enucleation of erythrocytes^15^, strongly support vasculogenesis^42^ and display numerous tissue-specific imprinted functions^43^. We have shown that lack of HLA class II expression is an intrinsic property of human primitive TRMs. This allows the distinction between primitive and definitive human macrophages, and can be harnessed to show that macrophage turnover dynamics in human fetal tissues correlate well with established dynamics in the mouse^5–8, 22^. However, we cannot exclude that the temporal increase in HLA class II expression in certain TRM populations (e.g. Kupffer cells and gut TRMs) might be due to an intrinsic upregulation in response to stimulation^44^. Functionally, the lack of HLA class II expression in primitive macrophages might simply reflect the lack of T cells when primitive haematopoiesis occurs^45, 46^. Microglia have minimal contributions from definitive haematopoiesis in mice^5, 6, 10^, yet express high levels of HLA class II in adult humans^47, 48^, suggesting that they either slowly acquire expression or that microglial turnover dynamics in humans are distinct from mice^49^.

Our findings demonstrate that the placenta is an additional site of primitive haematopoiesis in humans. Although CD34^+^ progenitors have been identified in first trimester placentas^15, 28, 29, 50^, they have been poorly defined. Placental primitive haematopoiesis is not supplanted by a transient definitive wave, as in the YS. Instead, PEMP maintain their ability to generate HLA-DR^neg^ primitive macrophages up until 8wk EGA, when their numbers begin to fall. The temporal distribution of PEMP currently remains unexplained. It is possible that PEMP have limited capacity for long-term self-renewal and so begin to disappear as pregnancy continues. Alternatively, an external cue might cause PEMP demise. A likely candidate is the sharp increase in oxygen tension and oxidative stress which occurs within placental villi between weeks 8 and 9 of gestation when the haemochorial circulation is established^51^.

Transcriptomic comparisons within and between species confirmed that PEMP are true EMPs, and that they lack the expression of *HLF*, a transcription factor constitutively expressed in definitive HSCs^38, 52^ and a critical regulator of HSC quiescence^53^. Although it cannot be ruled out that PEMP traffic to the placenta instead of being generated *in situ* we propose that this is unlikely as the placental vasculature does not mature until ∼10 wk GA^17, 18^. Furthermore, cells such as HLA-DR^pos^ monocytes, from the transient definitive wave of haematopoiesis that occurs in the yolk sac, are not found in the first trimester placenta^16^. Finally, we find PEMP in close association with developing placental endothelium and PEMP are transcriptionally and functionally distinct from YS progenitors (YSMP) found at equivalent developmental ages; as the latter express *HLF* and are poor at generating erythrocytes^20^.

In summary, our findings demonstrate that PEMP within early first trimester placental villi can give rise to HBC. Our establishment of *in vitro* assays to study PEMP provide a valuable tool to further develop our understanding of factors regulating primitive haematopoiesis and the generation of primitive macrophages in humans.

**Fig. S1.**
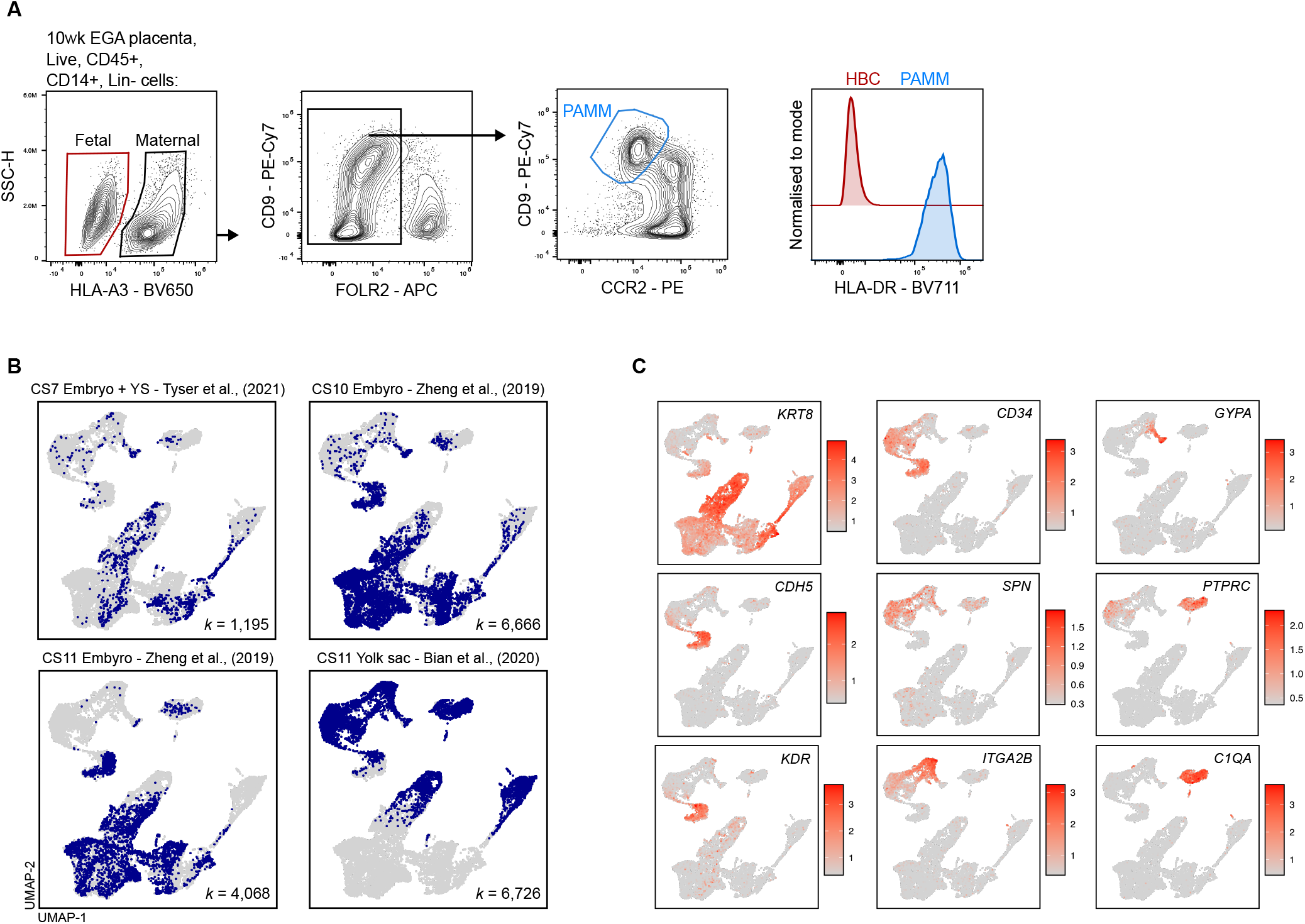
HBC HLA-DR expression and early human embryonic scRNAseq analysis. **A**) Representative flow cytometry gating strategy identifying fetal HBC (red), and maternal PAMM – Placenta-associated maternal macrophages (blue), and flow cytometry analysis of HLA-DR expression in HBC and PAMM. **B**) UMAP visualisation of human early embryo single-cell transcriptomes as in Fig.1A, with cells from each study highlighted in blue within each panel. Number of cells from each dataset (*k*) are annotated on each panel. **C**) UMAP visualisation with heatmap overlays of log-normalised expression of key marker genes for the identification of different lineages. *KRT8* – Non-heamatopoietic cells, *CDH5* and *KDR* – Endothelial cells, *CD34* and *SPN* – YSMP and haematopoietic progenitors, *ITGA2B* and *GYPA* – Megakaryocyte and Erythroid, *PTPRC* and *C1QA* – Macrophage.

**Fig. S2.**
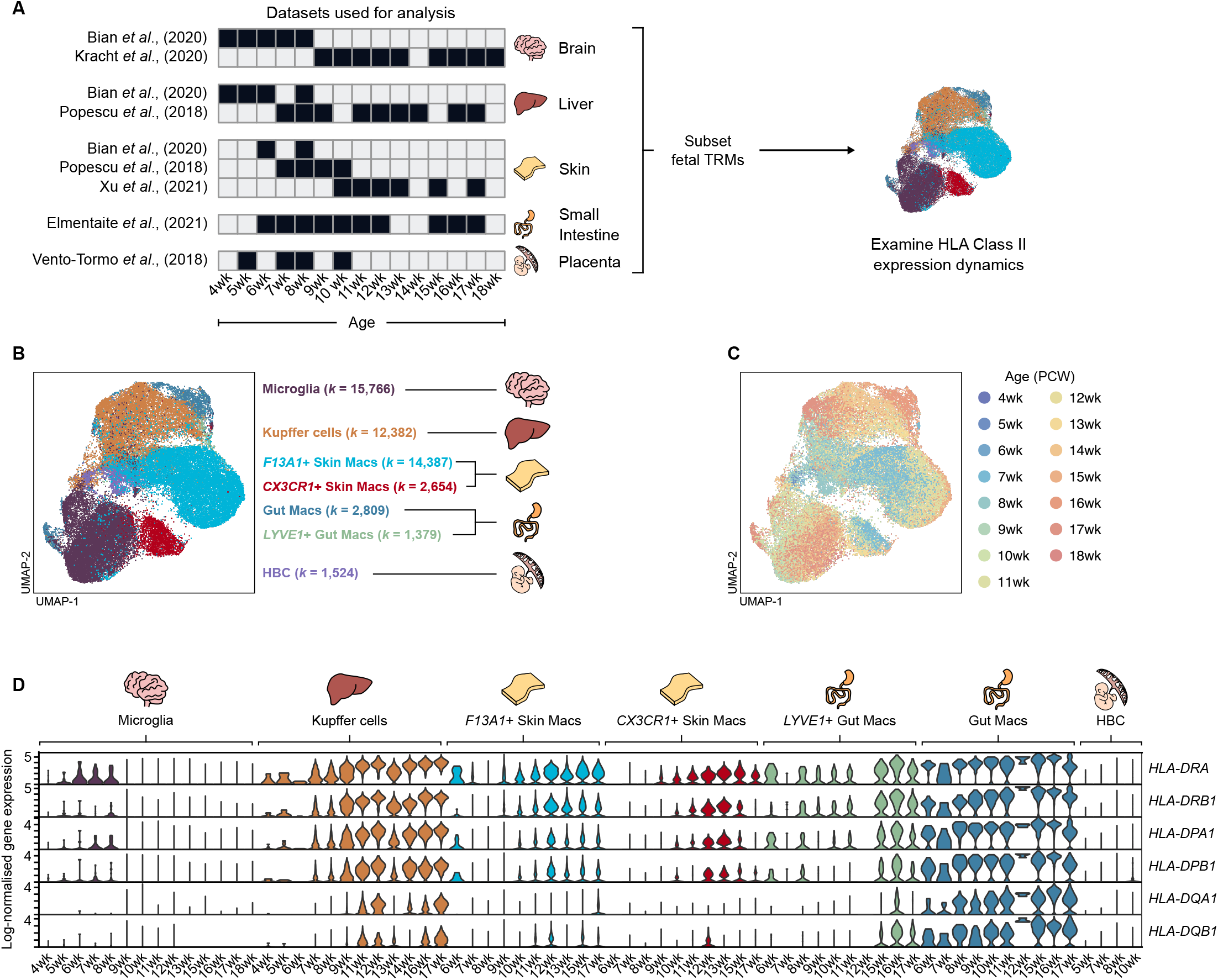
scRNAseq analysis of human TRMs during fetal development. **A**) Schematic representation of scRNAseq datasets used to construct human fetal TRM timelines from the brain^20, 27^, liver^20, 23^, skin^20, 23, 24^, small intestine^25^ and placenta^26^. Black squares denote cells from a given dataset and age were used. TRMs were subset and batch correction performed, correcting for the study of origin of all cells. **B**) UMAP visualisation of 50,901 human fetal tissue-resident macrophages (TRM) across developmental time, as in Fig1.E, with cells coloured by their identity. Datasets from^20, 23–27^. Number of cells of each identity (*k*) are annotated. **C**) UMAP visualisation with cells coloured by their developmental age in PCW – post-conception weeks. **D**) Log-normalised gene expression violin plots of HLA Class II genes in fetal TRMs across developmental time. The genes shown are used to calculate the proportion of cells which are HLA Class II^pos^ in Fig.1E.

**Fig. S3.**
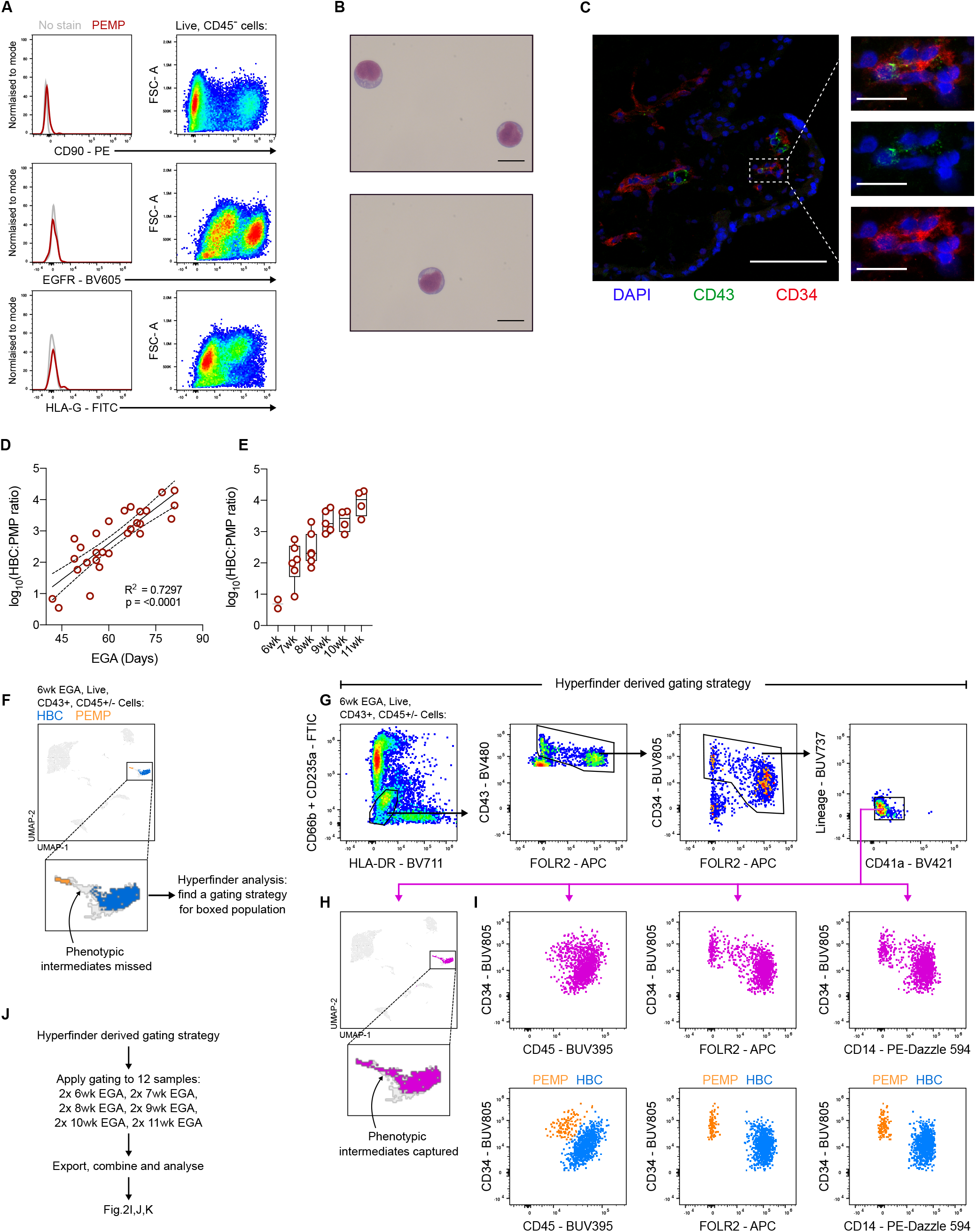
Phenotypic analysis of PEMP. **A**) Analysis of CD90, EGFR and HLA-G expression in PEMP and Live, CD45^-^ placental cells via flow cytometry, indicating that PEMP are not fibroblast or trophoblast contaminants. **B**) Raw Giemsa-stained cytospins of PEMP, as used in Fig.2I. **C**) Identification of CD34^+^ (red) CD43^+^ (green) PEMP within the villous stroma of a 7wk EGA placental sample. Scale bars, main panel 100μm, inset panels 20μm. **D**) Quantification of the ratio of HBC:PEMP determined by flow cytometry, plotted against EGA by day, or **E**) grouped by week. (*n* = 28). (F-I) Development of a gating strategy for the isolation of HBC, PEMP and any phenotypic intermediates. **F**) UMAP visualisation of flow cytometry data from live CD43^+^, CD45^+/-^ cells from a 6wk EGA placenta, with gated HBC (Blue) and PEMP (Orange) overlain. Phenotypic intermediates between HBC and PEMP within UMAP embedding are missed by conventional gating. **G**) Hyperfinder-derived gating strategy for the isolation of HBC, PEMP and any phenotypic intermediates, and **H**) resultant gated population overlain onto original UMAP in pink. **I**) Phenotypic analysis of cells obtained from Hyperfinder-derived gating strategy (pink) compared to typical PEMP (orange) and HBC (blue) gating, revealing the capture of phenotypic intermediates. **J**) Analysis workflow for the use of the Hyperfinder-derived gating strategy for the generation of Fig.2K-M.

**Fig. S4.**
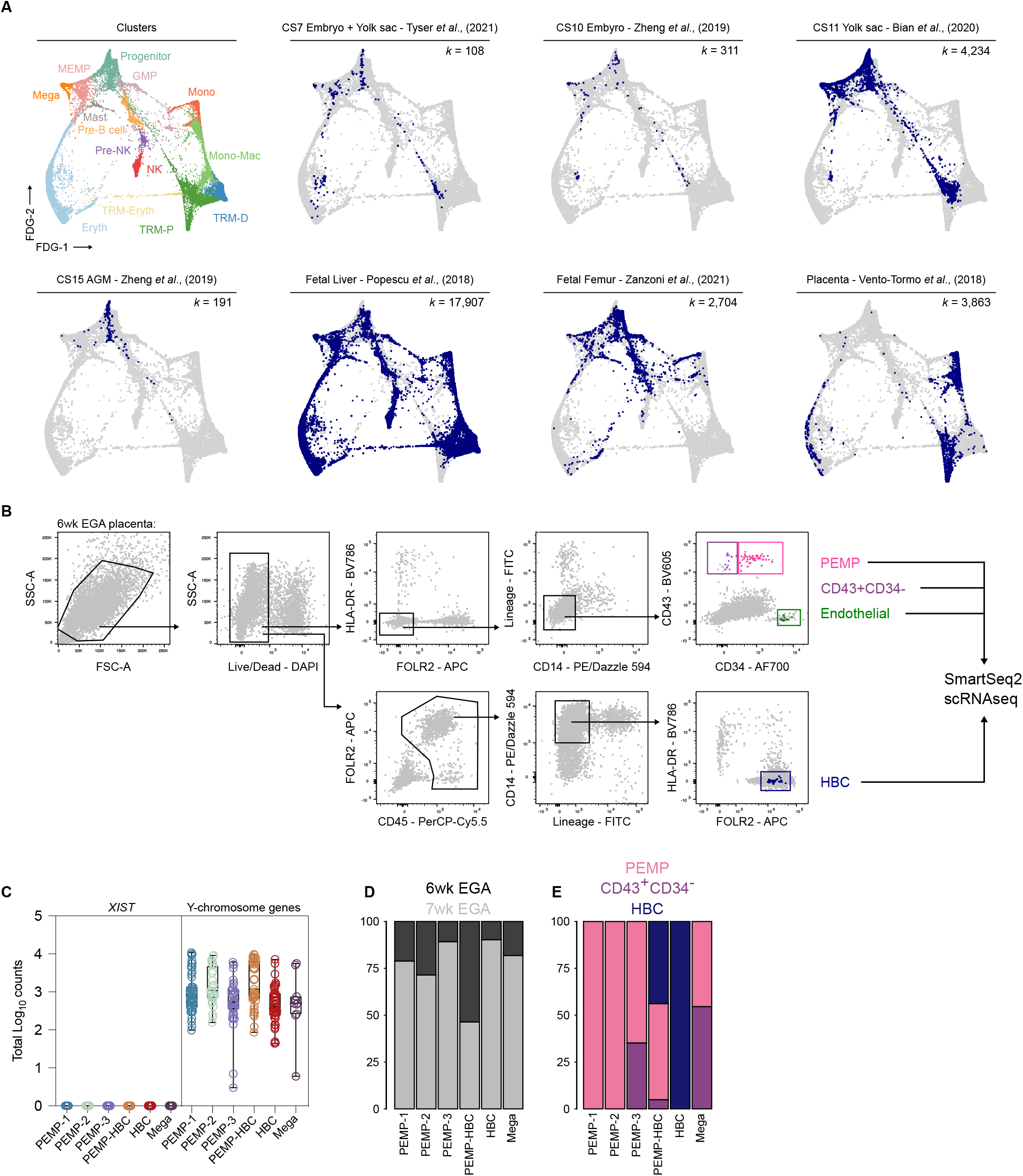
scRNAseq of PEMP from early placental samples. **A**) Force-directed graph embedding of 29,318 human haematopoietic cells from a range of anatomical sites and developmental timepoints. Data derived from^19–21, 23, 26, 33^. Top left panel – cells are coloured by their cluster identity. Other panels - cells from each study are highlighted in blue within each panel. Number of cells from each dataset (*k*) are annotated on each panel. GMP – Granulo-myeloid progenitor, MEMP – Megakaryocyte-Erythroid-Mast cell progenitor, Mega – Megakaryocyte progenitor, Eryth-Erythroid cells, Mono – Monocytes, Mono-Mac – Monocyte-derived macrophage, TRM-D – Definitive tissue-resident macrophages, TRM-P – Primitive tissue-resident macrophages, NK – Natural Killer cells. **B**) Representative FACS gating strategy for the isolation of PEMP, CD34^+^ CD43^+^, endothelial cells and HBC for plate-based SmartSeq2 scRNAseq. **C**) Boxplots showing the log-normalised total counts for *XIST* and for all genes from the Y-chromosome for each cluster in the scRNAseq dataset. XIST expression was not detected in any cell in the dataset, and all cells displayed Y-chromosome-specific counts, suggesting no contamination of maternal cells, and that all profiled cells are fetal in origin. (D-E) Stacked bar charts indicating the proportion of cells from each cluster **D**) derived from each donor or **E**) derived from each gate during FACS isolation.

**Fig. S5.**
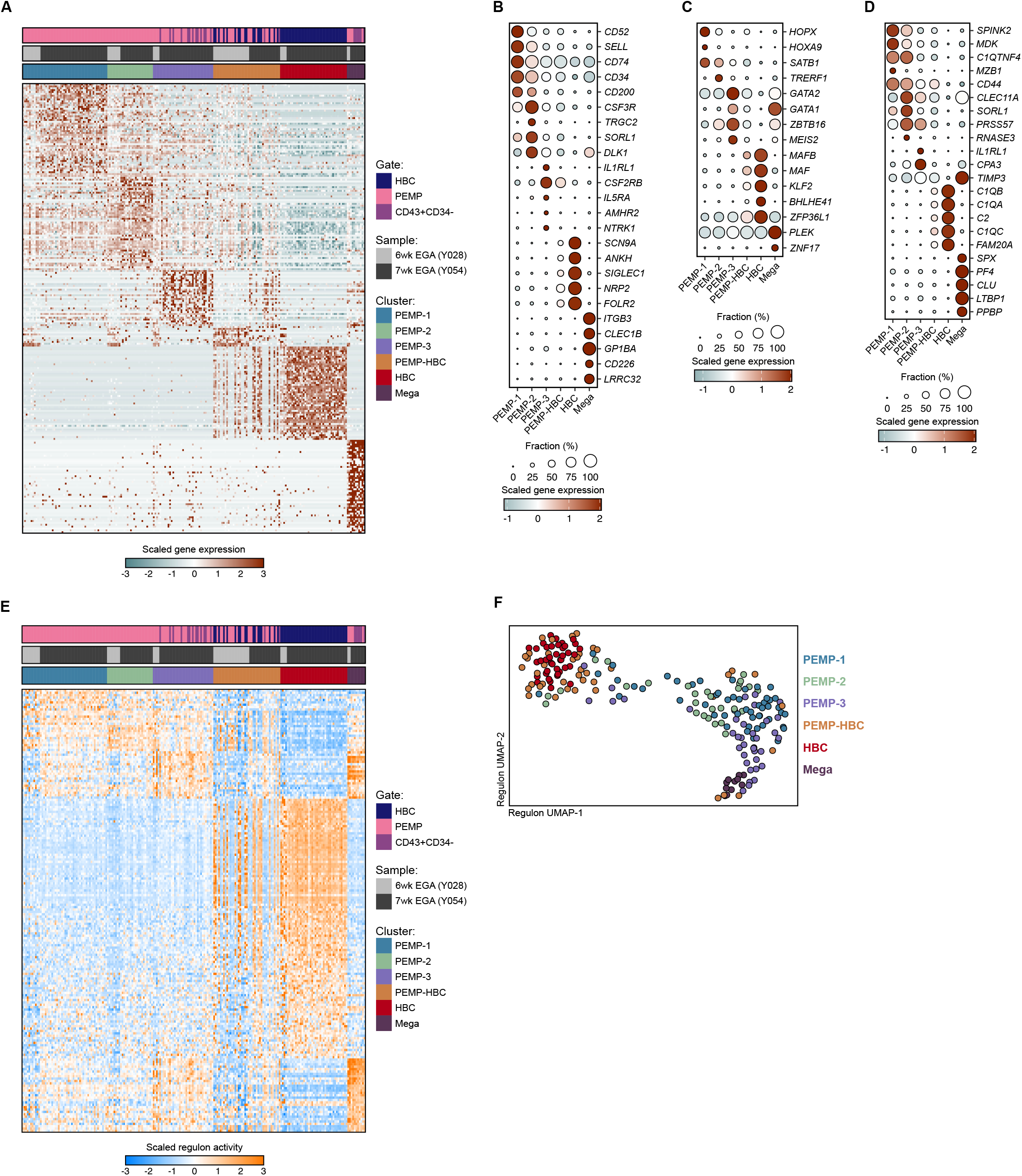
Differential gene expression and transcription factor activity across scRNAseq clusters. **A**) Heatmap showing scaled log-normalised gene expression of the top 50 differentially expressed genes per cluster from the scRNAseq dataset. (B-D) Dotplot heatmap displaying scaled log-normalised gene expression of the top 5 **B**) surface markers **C**) transcription factors and **D**) secreted factors per scRNAseq cluster. Dot size represents fraction of cells with nonzero expression. **E**) Heatmap showing scaled inferred transcription factor activity (regulon scores) of all regulons with differential activity across clusters from the scRNAseq dataset. **F**) UMAP visualisation of placental scRNAseq dataset with embeddings calculated using regulon scores instead of gene expression data.

**Fig. S6.**
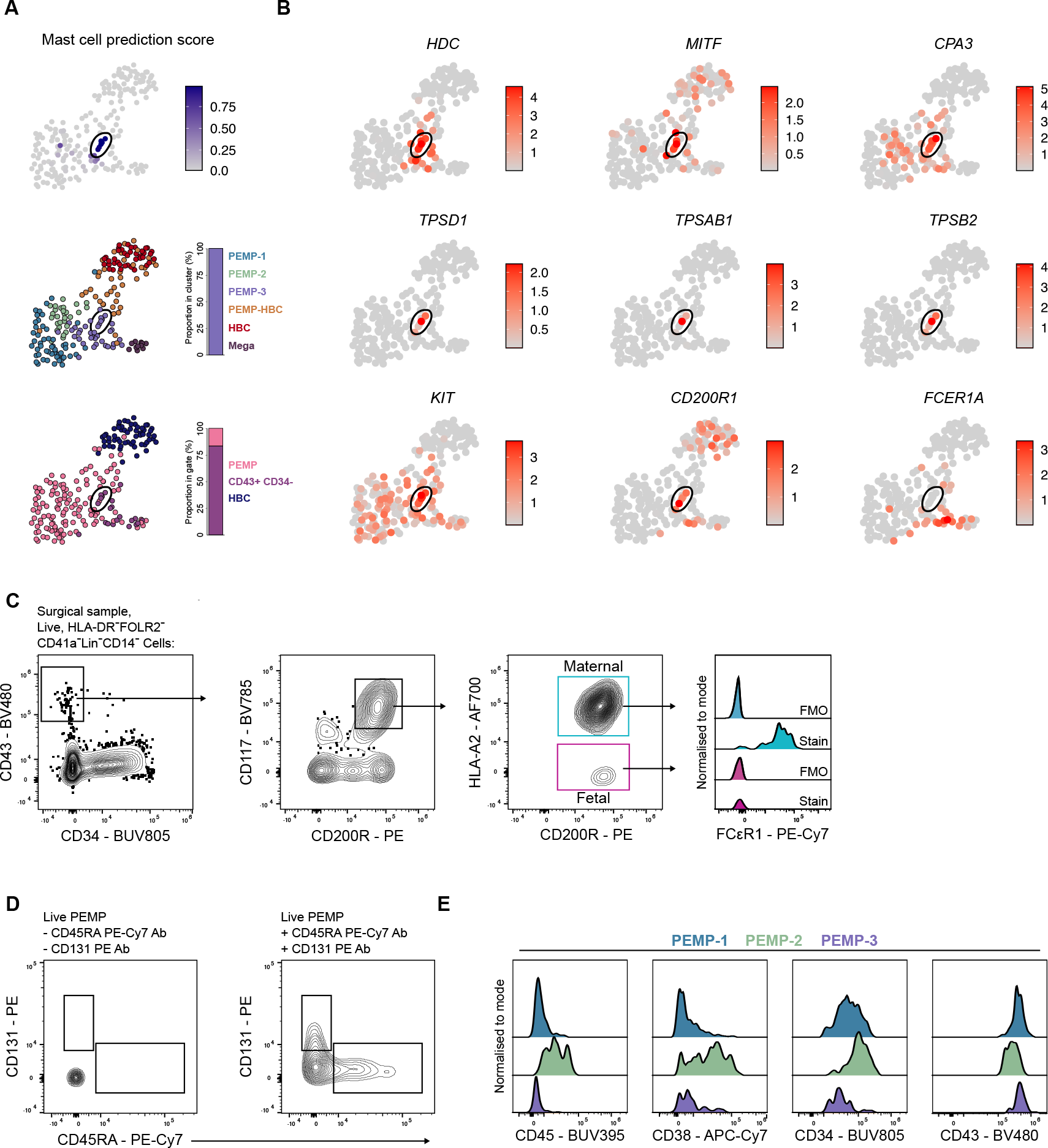
Fetal placental mast cells and PEMP phenotypic heterogeneity. **A**) UMAP visualisation of placental scRNAseq dataset with fetal liver mast cell prediction score from Fig.3D overlain. Cells with high mast cell prediction scores are circled, and the proportions of these cells in each cluster and FACS gate are indicated in the stacked bar charts in the lower panels. **B**) UMAP visualisation of placental scRNAseq dataset with overlays of log-normalised expression of mast cell-specific genes. Cells with high mast cell prediction scores are circled. Fetal placental mast cells lack the expression of *FCER1A* consistent with previous findings^35, 36^. **C**) Identification of maternal CD200R^+^CD117^+^FCεR1^+^ and fetal CD200R^+^CD117^+^ FCεR1^-^ mast cells from a first trimester placenta surgical sample via flow cytometry. Maternal mast cells are likely derived from contamination of the maternal decidua, which is more prevalent in surgical samples. FMO – Fluorescence minus one. **D**) Analysis of CD131 and CD45RA expression within PEMP via flow cytometry without CD131 and CD45RA antibodies added (left panel) and with antibodies added (right panel). **E**) Analysis of CD45, CD38, CD34 and CD43 expression in ^-/-^ PEMP (blue), CD45RA^+^ PEMP (green) and CD131^+^ PEMP (purple) via flow cytometry. Data shown are representative of n= 4 donors.

**Fig. S7.**
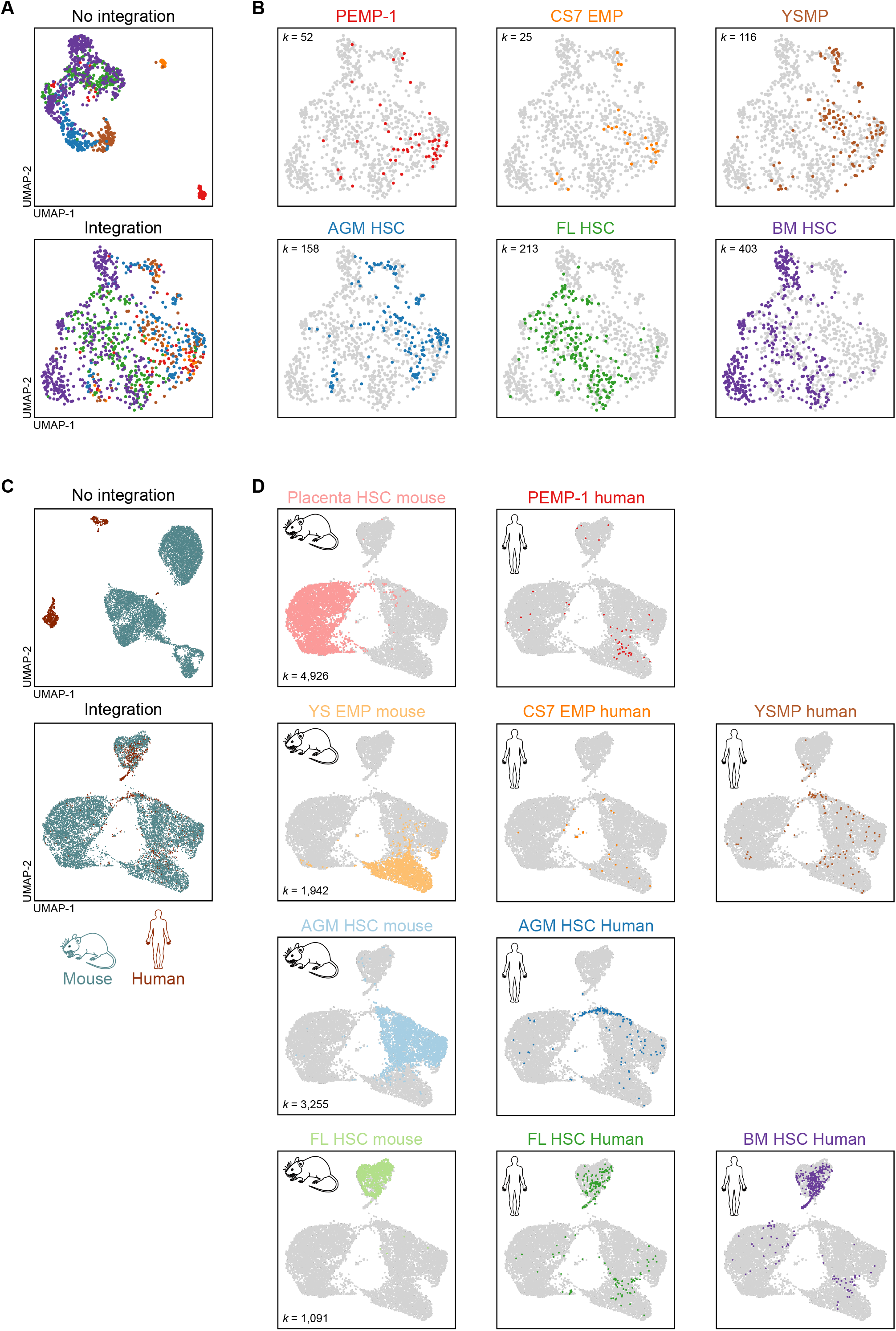
Intra- and inter-species fetal haematopoietic progenitor datasets. **A**) UMAP visualisations of 967 single cells from human haematopoietic progenitor populations used for comparative analysis (Fig.4B)^19–21, 33^, shown before (upper panel) and after (lower panel) Seurat V3 integration^54^, correcting for the study of origin of each cell. Cells are coloured according to identity, see panel B. **B**) UMAP visualisation of cells of each identity in individual panels. Number of cells of each identity (*k*) are annotated on each panel. CS – Carnegie Stage, EMP – Erythro-myeloid progenitors, YSMP – Yolk sac myeloid-biased progenitors, AGM – Aorta-gonad mesonephros, FL – Fetal liver, BM – Bone marrow, HSC – Haematopoietic stem cell. **C**) UMAP visualisations of 12,181 human and murine fetal haematopoietic progenitors used for comparative analysis^19–21, 33, 39, 40^ shown before (upper panel) and after (lower panel) Seurat V3 integration^54^, correcting for the species of each cell. Cells are coloured according to their species. Mouse data were “humanised” for this analysis by replacing murine genes with direct human homologs (see methods). **D**) UMAP visualisation of cells of each identity in individual panels. Number of murine cells of each identity (*k*) are annotated on each panel. YS – Yolk sac.

**Fig. S8.**
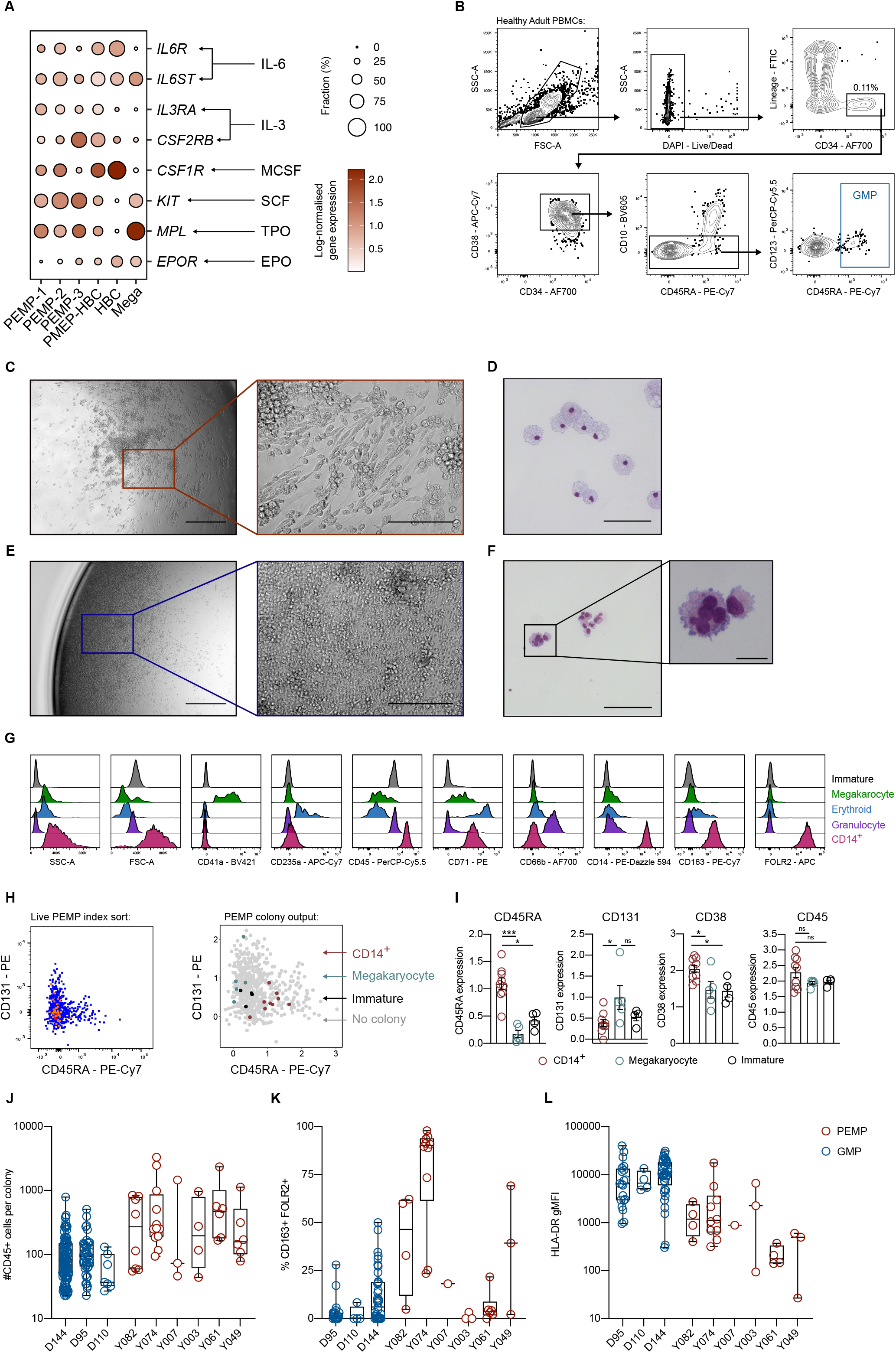
PEMP single-cell differentiation assays. **A**) Dotplot heatmap displaying log-normalised gene expression of cytokine receptor genes within the placental scRNAseq dataset. Cognate ligands which are components of the PEMP differentiation cytokine cocktail are annotated onto the plot with arrows. Dot size represents fraction of cells with nonzero expression. **B**) Representative FACS gating strategy for the isolation of healthy adult blood GMP (Granulo-myeloid progenitors) for single cell differentiation assays. Data representative of n = 3 donors. **C**) Representative morphology of a CD14^+^ colony derived from a single PEMP after 18 days of culture on primary placental fibroblasts. Scale bars, 500μm left panel, 100μm right panel. Representative image from n=6 donors. **D**) Representative Giemsa-stained cytospins of cells from a CD14^+^ colony derived from a single PEMP. Scale bars, 100μm. Representative image from n=4 donors. **E**) Representative morphology of a diverse haematopoietic colony derived from a single PEMP after 18 days of culture on primary placental fibroblasts. Scale bars, 500μm left panel, 100μm right panel. Representative image from n=6 donors. **F**) Representative Giemsa-stained cytospins of cells from a diverse haematopoietic colony derived from a single PEMP. Inset, morphology of a suspected PEMP-derived polyploid megakaryocyte. The colony from which this cytospin was taken was found to have made megakaryocytes via flow cytometry analysis. Scale bars, 100μm left panel, 20μm right panel. Representative image from n=4 donors. **G**) Comparative analysis of marker expression via flow cytometry in distinct immune lineages derived from PEMP. **H**) Flow cytometry analysis of CD45RA and CD131 expression in 576 index-sorted PEMP from *n*=2 donors (left panel), and colony output from resulting cultures (see Fig.5.D,E for colony gating strategy) overlain onto logicle-transformed biaxial plot of CD45RA and CD131 expression. Annotated colonies are gated as in Fig.5D,E: CD14^+^ – Live, CD45^+^CD14^+^CD66b^-^, Megakaryocyte – Live, CD41a^+^, Immature – Live, CD45^+^. **I**) Quantification of the expression of CD45RA, CD131, CD38 and CD45 by index-sorted PEMP which give rise to macrophage, megakaryocyte and immature colonies. P-values calculated by Mann-Whitney test. (H-J) Boxplot quantifications of **H**) number of CD45^+^ cells per colony, **I**) percentage co-expression of CD163 and FOLR2 expression in CD14^+^ colonies and **J**) HLA-DR expression in CD14+ colonies, for each donor profiled. NS, P > 0.05, *, P ≤ 0.05, ***, P ≤ 0.001.

## Methods and Materials

### Patient samples

All tissue samples used were obtained with written consent from participants. Placental tissues were obtained from healthy women with apparently normal pregnancies undergoing elective first trimester terminations (6-12 weeks EGA). The EGA of the samples was determined from the last menstrual period. Peripheral blood was taken from healthy adult volunteers. All samples were obtained with written informed consent from participants under ethical approval which was obtained from the Cambridge Research Ethics committee (study 04/Q0108/23).

### Tissue processing

Placental samples were processed immediately upon receipt as previously reported^16^. Samples were washed in PBS for 10 minutes with a stirrer before processing. The placental villi were scraped from the chorionic membrane with a scalpel and digested with 0.2% Trypsin (Pan-Biotech)/ 0.02% Ethylenediaminetetraacetic acid (EDTA) (Source BioScience) at 37°C with stirring, for 7 minutes. The digested cell suspension was passed through a sterile muslin gauze, and fetal bovine serum (FBS) (Sigma-aldrich) was added to halt the digestion process. The undigested tissue left on the gauze was scraped off with a scalpel and digested in 2.5ml 1mg/ml collagenase V (Sigma-Aldrich), supplemented with 50μl of 10mg/ml DNAse I (Roche) for 20 minutes at 37°C with agitation. The digested cell suspension was passed through a sterile muslin gauze and washed through with PBS. Cell suspensions from both the trypsin and collagenase digests were pelleted, resuspended in PBS and combined. Cells were layered onto a Pancoll gradient (PAN-biotech) and spun for 20 minutes without brake at 3,000 rotations per minute (rpm). The leukocyte layer was collected and washed in PBS. Blood samples were processed as described previously^16, 55^.

### Flow cytometry and data analysis

Cell suspensions were stained for viability with 1:1000 LIVE/DEAD Fixable Blue (Thermo Fisher Scientific), or 1:1000 Zombie Aqua (Biolegend), both for 20 minutes at 4°C, and washed twice in PBS. For cell sorting cell suspensions were stained for viability with 1:3000 4′,6-diamidino-2-phenylindole (DAPI) (Sigma-Aldrich) immediately before sorting.

Cells were blocked in human blocking buffer (5% human serum (Sigma-Aldrich), 1% rat serum (Sigma-Aldrich), 1% mouse serum (Sigma-Aldrich), 5% FBS and 2mM EDTA) for 15 minutes at 4°C and were incubated with antibody cocktails for 30 minutes at 4°C. Antibodies used are listed in **Supplementary Table 1**. Cells were washed and resuspended in FACS buffer (PBS containing 2% FBS and 2mM EDTA). The lineage (lin) channel in flow cytometry analyses included combinations of the markers CD3, CD19, CD20, CD41, CD56, CD66b, CD235a and CD335, for the removal of contaminating T cells, B cells, NK cells erythrocytes, megakaryocytes/platelets and granulocytes. Flow cytometry was performed using a Cytek Aurora (Cytek) or an Attune NxT (Thermo Fisher Scientific), or cells were purified by cell-sorting using a BD FACS Aria III (BD bioscience). Flow cytometry data was analysed using FlowJo v10.7 (Treestar), R version3.6.3 (The R foundation) and Prism 9 (GraphPad).

A gating strategy for the isolation of PEMP, HBC and any intermediates was identified using the Hyperfinder plugin in Flowjo, using default parameters. Down-sampled gated cells were exported from Flowjo, imported into R and transformed using the ‘*autoLgcl’* transformation using the ‘*cytof_exprsMerge’* function from the Cytofkit2 R package. The datasets were batch corrected using the ‘*mnnCorrect’* function from the batchelor R package and subjected to PCA analysis using the ‘*prcomp’* function.

Index-sort plate data was loaded into R using the ‘*retrieve_index’* function from the indexSort R package, and the resultant data matrices were transformed via the ‘*logicleTransform’* function in the flowCore R package (parameters; *w* = 0.6, *m* =4.2, *a* = 0). Cells which generated colonies were manually selected and transformed marker expression data was extracted for visualisation and statistical analysis in Prism 9 (GraphPad).

### Cytospins of sort-purified cells

PEMP were sorted by FACS as detailed above. Cells from PEMP-derived colonies were taken for cytospins before they were stained for flow cytometry. Cells were resuspended in 150μl of PBS and centrifuged onto glass slides using a Shandon Cytospin 2 (Marshall Scientific) at 300 rpm for 5 minutes. Cells were fixed in methanol for 2 minute and air-dried. Slides were stained with 1:30 Giemsa-stain (Sigma-Aldrich) for 25 minutes and mounted in DePex mounting medium (BHD). Slides were imaged under a 63X objective on a Zeiss AxioObserver Z1 Microscope (Zeiss).

### Immunofluorescence of placental tissue sections

Sections from early EGA first trimester placenta villous tissue were prepared as described previously. Slides were placed in blocking buffer for 20 minutes at room temperature, washed in PBS and incubated overnight at 4°C with primary antibodies (**Supplementary Table 1**). The slides were washed twice for 5 minutes in PBS, incubated with secondary antibodies for 1h at room temperature and washed twice for 5 minutes in PBS. The slides were then stained with 1/1000 4′,6-diamidino-2-phenylindole (DAPI) (Sigma-Aldrich) for 10 minutes at room temperature and washed twice for 5 minutes in PBS. Slides were mounted using ibidi mounting medium (ibidi). Slides were imaged using a Zeiss SP8 confocal LSM 700 (Zeiss).

### Single cell differentiation assays

Primary placental fibroblasts were obtained by culturing placental digests in Dulbecco’s Modified Eagle Medium (DMEM) (Thermo Fisher Scientific) supplemented with 10% FBS, 2.5% Penicillin Streptomycin (Pen/Strep) (Sigma-Aldrich) and 20μM L-Glutamine (Sigma-Aldrich). Non-adherent cells were removed, and cells were passaged a minimum of 4 times, yielding a pure population of fibroblasts, which were cryopreserved. Placental fibroblasts were plated at a density of 3,000 cells/well into flat 96-well plates in 100ul DMEM (Thermo Fisher Scientific) supplemented with 10% FBS, 2.5% Pen/Strep (Sigma-Aldrich) and 20μM L-Glutamine (Sigma-Aldrich) 2-3 days before sorting. On the day of the sort, the medium was changed to 100μl/well StemPro-34 medium with nutrient supplement (Thermo Fisher Scientific) supplemented with cytokines (IL-6 10ng/ml, IL-3 20ng/ml, M-CSF 50ng/ml and SCF 20ng/ml; all Miltenyi Biotec), with 2.5% Pen/Strep and 20μM L-Glutamine (Sigma-Aldrich). Cryopreserved placental and adult peripheral blood cells were used for single-cell differentiation assays, cells were thawed, washed, and stained for FACS as detailed above. Cells were index sorted (1 cell/well) and cultured for a total of 18 days. On day 7 an additional 50μl of media with cytokines was added, with the addition of EPO (Expex) and TPO (Miltenyi Biotec) to final concentrations of 1.5 units\ml and 25ng/ml respectively. EPO and TPO were not added until day 7 to favour the differentiation of macrophages. Brightfield images of colonies were taken on an EVOS M5000 microscope (Thermo Fisher Scientific). After 18 days plates containing colonies were incubated on ice for 15 minutes and colonies were removed from culture plates via gentle aspiration and washed in PBS. Colonies were then analysed by flow cytometry or cytospin as detailed above. Lineage potentials were ascertained when the numbers of cells of the indicated phenotypes were more than 30 (PEMP) or 20 (GMP) per well.

### SmartSeq2 scRNAseq of placental cells

Live cells were isolated by FACS as detailed above and sorted into 96 well plates containing 2.3μl of SmartSeq2 lysis buffer^56^ per well (10% SUPERase-ln RNase inhibitor (Thermo Fisher Scientific), 0.2% Triton X-100 (Sigma-Aldrich). Once cells were collected plates were sealed, spun at 700g for 1 minute and frozen using dry ice before storage at −80°C. Sequencing libraries were constructed using the SmartSeq2 protocol^56^ and 150 BP paired-end sequencing was performed using a Illumina Novaseq SP.

### SmartSeq2 scRNAseq data pre-processing and visualisation

SmartSeq2 sequencing data were trimmed with Trim Galore (version 0.4.0, https://github.com/FelixKrueger/TrimGalore) default settings, low quality (Q<20) and short reads (<20bp) were removed. Remaining reads were then aligned with STAR (v 2.5.0.a)^57^ using the GRCh38 human reference genome and annotation, supplemented with External RNA Controls Consortium (ERCC) spike-in controls‘ERCC92.fasta’ (https://gist.github.com/mikheyev/821f96a0ce76f58ee0ba). Gene-specific read counts were calculated using ‘featureCounts’ function in subread package (version 1.6.2)^58^.

Downstream analyses of gene expression matrices were performed using Seurat (v3.0)^54^. Cells with fewer than 3,500 (Y028) or 2,000 (Y054) detected genes, and more than 30% mitochondrial gene expression were removed, yielding a total of 255 cells. Samples were log-normalised and pre-processed individually following the Seurat v3 workflow. Clusters were identified in each dataset using the *FindNeighbours* and *FindClusters* functions in Seurat. Clusters were annotated on the basis of expression of known marker genes. For the purposes of this study 35 endothelial cells (*CDH5*+, *KDR*+) and 10 erythrocytes (*HBZ*+, *HBE1*+) were removed from the datasets. The two samples were merged and integrated via the Seurat v3 integration workflow. Clusters were identified using the *FindNeighbours* and *FindClusters* functions in Seurat. Uniform Manifold Approximation and Projection (UMAP) dimensionality reduction was performed using the *RunUMAP* function in Seurat, using the first 20 principal components (PCs). Significantly differentially expressed genes (DEGs) were identified using the *FindMarkers* function, using the Wilcox rank sum test, corrected for multiple comparisons.

### Analysis of publicly available scRNAseq data and generation of combined fetal scRNAseq datasets

Details and sources of publicly available datasets used in this study can be found in **Supplementary Table 1** All datasets were processed as detailed above using Seurat. Datasets available as ‘.*h5ad’* format were converted into Seurat objects using the ‘*convert’* and ‘*LoadH5Seurat’* functions in the R package SeuratDisk. Where available, cluster annotations from the original studies were added to the meta-data of each object to allow for the identification of distinct cell types. Where annotations were not available clusters were manually annotated based on known marker genes, using the source publications as references.

For the construction of the early human embryo object (**Fig.1A**) datasets from a CS7 Embryo (E-MTAB-9388, http://www.human-gastrula.net/)^19^, CS10 embro body (GSE135202)^21^, CS11 Caudal half (GSE135202)^21^ and CS11 yolk sac (GSE137010)^20^ were merged, and batch correction performed using the ‘*RunFastMNN’* function in the SeuratWrappers R package, correcting for the tissue of origin. Clustering and dimensionality reduction were performed using the first 20 MNN components, and clusters were annotated based on known marker gene expression.

The human fetal TRM dataset (**Fig.1E, F**, **S2A-D**) comprised data from early human embryos (GSE133345)^20^, fetal microglia (GSE141862)^27^, fetal liver (E-MTAB-7407)^23^, fetal skin (E-MTAB-7407 and GSE179565)^23, 24^, fetal gut (E-MTAB-8901, https://www.gutcellatlas.org/)^25^ and placenta (E-MTAB-6701)^26^. Each dataset was subset to include only TRMs. The early human embryo dataset was first subset by tissue of origin (head, liver or skin), and then subset to only include TRMs for each organ. As annotations were not available for the fetal skin datasets (GSE133345, E-MTAB-7407 and GSE179565), the original datasets were merged, batch-corrected and two major populations of *F13A1*+ and *CX3CR1*+ TRMs were identified via clustering and dimensionality reduction, consistent with source publications. All TRM datasets were then merged, and batch correction performed using the ‘*RunFastMNN’* function correcting for the study each dataset was derived from. Dimensionality reduction was performed using the first 10 MNN components and TRMs were annotated based on both their identity and age for gene expression analysis. The proportion of TRMs expressing HLA Class II (Fig.1E) was determined by calculating the proportion of cells co-expressing *HLA-DRA*, *HLA-DRB1*, *HLA-DPA1*, *HLA-DPB1*, *HLA-DQA1* and *HLA-DQB1.* Co-expression was defined as a cell exhibiting non-zero expression of all genes simultaneously.

The human fetal haematopoiesis dataset (**Fig.S4A**) comprised data from a CS7 Embryo (http://www.human-gastrula.net/)^19^, CS10 embryo body (GSE135202)^21^, CS11 yolk sac (GSE137010)^20^, CS15 AGM (GSE135202)^21^, Fetal liver (E-MTAB-7407, https://www.covid19cellatlas.org/index.healthy.html)^23^, Fetal Femur (E-MTAB-9067, https://gitlab.com/cvejic-group/integrative-scrna-scatac-human-foetal/-/tree/master/Data/ScanpyObjets)^33^ and placenta (E-MTAB-6701)^26^. The fetal liver dataset was downsampled from 95,461 cells to 20,000 cells (maintaining the structure of the data) to ease computational burden. Datasets were merged, and batch correction performed using the ‘*RunFastMNN’* function correcting for the tissue of origin. From the resultant object haematopoietic cells were identified, subset and subjected to an additional round of batch correction, yielding the final object shown in the figure. A force-directed graph embedding was computed using the first 20 MNN components using ForceAtlas2 with 600 iterations.

### SCENIC analysis

To infer transcription factor activity in our scRNAseq data, gene regulatory network analysis was performed with SCENIC^34^. Regulons were inferred following the SCENIC analysis pipeline, and regulon activity scores for each cell were added as a new *Assay* to the scRNAseq Seurat object, permitting downstream analysis. Regulons with differential activity between clusters were calculated using *FindMarkers* function, and selected regulon activities were plotted using *pheatmap.* The regulonUMAP embedding was computed using regulon activity scores in the SCENIC-regulon assay of the Seurat object, with the first 15 principle components used.

### Comparisons of placental cells with an atlas of fetal liver haematopoiesis

Fetal liver haematopoiesis and placental scRNAseq datasets were processed as detailed above. The *FindTransferAnchors* function in Seurat was used to compare the transcriptomes of the query dataset (placental) with the reference dataset (Fetal liver) using the top 30 PCs. Prediction scores for each cell in the placental dataset were calculated for each Fetal liver cluster, with higher prediction scores indicating a higher level of transcriptional similarity between a given query cell and a given reference cluster. Mean prediction scores for each placental scRNAseq cluster were plotted as a heatmap using *pheatmap*.

### Comparisons of PEMP-1 with other human fetal haematopoietic progenitors

Each individual progenitor population was subset from its original dataset; ‘CS7 EMP’ from CS7 Embryo (E-MTAB-9388, http://www.human-gastrula.net/)^19^, ‘YSMP’ from CS11-17 YS (GSE133345)^20^, ‘AGM HSC’ from CS15 AGM (GSE135202)^21^, ‘FL HSC’ and ‘BM HSC’ from fetal liver and fetal bone marrow (E-MTAB-9067, https://gitlab.com/cvejic-group/integrative-scrna-scatac-human-foetal/-/tree/master/Data/ScanpyObjets)^33^ and PEMP-1 from the scRNAseq dataset generated in this study. The datasets were merged, and integrated via the Seurat V3 integration workflow, correcting for the study of origin (**Fig.S7A,B**). Transcriptomic similarity between progenitor populations was assessed using the ‘*BuildClusterTree’* function in Seurat utilising the 2,000 features used for integration (**Fig.4B**). For DEG analysis the object was down-sampled to 50 cells per population to prevent cell numbers confounding the analysis.

DEGs were identified between CS7 EMP and definitive progenitors, and PEMP-1 and definitive progenitors using the *FindMarkers* function with *logfc.threshold* = 0, using the Wilcox rank sum test, corrected for multiple comparisons. DEGs with a logFC >0.5 or < −0.5 and an adjusted p-value of <0.05 in either or both analyses were visualised.

### Comparisons of PEMP-1 with murine fetal haematopoietic progenitors

Murine placental data was obtained from GSE152903^39^ and murine FL, AGM ad YS data was obtained from GSE137116^40^. To allow for the direct comparison of murine and human datasets we identified human homologs of murine genes from the gene expression count matrices, using the ‘*getLDS*’ function in the biomaRt R package. The murine count matrices were then subset to include only genes with direct human homologs, and murine gene annotations were replaced with the corresponding human gene annotation, in order to generate “humanised” murine scRNAseq data. “Humanised” murine data was pre-processed as detailed above and progenitor populations were identified either by manual data inspection (placental data) or provided annotations (FL, AGM ad YS data) and subset. Murine and human datasets were merged, and integrated via the Seurat v3 integration workflow, correcting for the species of origin (**Fig.S7C,D**). Transcriptomic similarity between progenitor populations was assessed using the ‘*BuildClusterTree’* function in Seurat utilising the 2,000 features used for integration (**Fig.4E**). DEGs were identified between PEMP-1 and murine placental HSC using the *FindMarkers* function. DEGs were also calculated between all human and murine progenitor populations (all down-sampled to 50 cells) to allow for the identification of DEGs which are independent of the difference in species.

## Acknowledgements

We thank the following for assistance: The Flow Cytometry Core Facility at the Department of Pathology, Cambridge. David C. J. Carpentier at the Microscopy Core at the Department of Pathology, Cambridge. Lucy Gardner for her help in processing placental samples. We thank all donors who participated in this study and hospital staff. Graham Burton for scientific discussion.

This work was supported by the Wellcome Trust, Royal Society, Centre for Trophoblast Research, and Department of Pathology, University of Cambridge, UK. N.McG, is funded by a Wellcome Sir Henry Dale and Royal Society Fellowship (grant number 204464/Z/16/Z). J.R.T is funded by a Wellcome Trust PhD Studentship (grant number 215226/Z/19/Z). AS is funded by the MRC (grant number: MR/P001092/1). E.L. is supported by a Sir Henry Dale fellowship from Wellcome/Royal Society (107630/Z/15/Z). Research in E.L.’s laboratory is supported by Wellcome (107630/Z/15/Z; 215116/Z/18/Z), BBSRC (BB/P002293/1), Royal Society and by core support grants by Wellcome and MRC to the Wellcome-MRC Cambridge Stem Cell Institute (203151/Z/16/Z). This research was funded in whole, or in part, by the Wellcome Trust. For the purpose of Open Access, the author has applied a CC BY 4.0 public copyright license to any Author Accepted Manuscript version arising from this submission.

## Data and Code availability

Raw and processed scRNAseq data will be published in GEO prior to publication. Any additional information required to reanalyse the data or reproduce any analyses in this paper will be available from the lead contact upon request.

## Author Contributions

Conceptualisation: J.R.T and N.McG. Methodology: J.R.T, A.A, E.FC, X.Z, R.SH, E.L. and NMcG. Formal analysis: J.R.T and N.McG. Intellectual input: J.R.T, X.Z, E.FC, R.SH A.M, A.S, E.L. and NMcG. Writing: J.R.T. and N.McG. Visualisation: J.R.T. and N.McG. Supervision: N.McG. All authors revised and discussed the manuscript.

